# A head-mounted multi-camera system for electrophysiology and behavior in freely-moving mice

**DOI:** 10.1101/2020.06.30.181412

**Authors:** Nicholas Sattler, Michael Wehr

## Abstract

Advances in the ability to monitor freely-moving mice may prove valuable for the study of behavior and its neural correlates. Here we describe a head-mounted multi-camera system for mice, comprised of inexpensive miniature analog camera modules. We illustrate the use of this system with several natural behaviors including prey capture, courtship, jumping, and exploration. With a four-camera headset, monitoring the eyes, ears, whiskers, rhinarium, and binocular visual field can all be achieved simultaneously with high-density electrophysiology. With appropriate focus and positioning, all eye movements can be captured, including cyclotorsion. For studies of vision and eye movements, cyclotorsion provides the final degree of freedom required to reconstruct the visual scene in retinotopic coordinates or to investigate the vestibulo-ocular reflex in mice. Altogether, this system allows for comprehensive measurement of freely-moving mouse behavior, enabling a more holistic and multimodal approach to investigate ethological behaviors and other processes of active perception.

## Introduction

Mice move. The whiskers whisk, the eyes saccade, the nostrils sniff, and the ears pivot (Preyer 1882; Welker 1964; Carvell et al. 1991; Wallace et al. 2013). These active sensing behaviors have been well-established as adaptive strategies for the optimization of somatosensation (Gibson, 1962; Lederman and Klatzky, 1987; Carvell and Simons, 1995; Bagdasarian et al., 2013), vision (Ballard, 1991; O'Regan and Noë, 2001), olfaction (Mainland and Sobel, 2006), and audition (Populin and Yin, 1998; Holland and Waters, 2005; Tollin et al., 2009). Recent studies have also revealed that movement has widespread neural correlates throughout the brain. Eye movements, whisking, sniffing, running, and other classes of movements can each produce independent effects on brain regions far removed from the directly-involved motor or sensory systems (Musall et al., 2019; Stringer et al., 2019; Salkoff et al., 2020). Investigating sensory processing therefore requires either that all movements are eliminated (e.g. by head or eye fixation), or that they are accurately measured. Despite the insights that have been gained from research under head-fixed or eye-fixed conditions, there has been a growing appreciation that understanding brain function will require recording neural activity in freely-moving animals, especially during natural behavior (Datta et al. 2019; Parker et al. 2020). Yet monitoring the simultaneous movements of the eyes, whiskers, nose, ears, limbs, and the rest of the body during freely-moving behavior and with high-density electrophysiology remains methodologically challenging.

There are many inherent technical difficulties when designing attached devices for freely-moving behavior in small animals. Every component comes with an associated cost of size, weight, positioning, balance, and cabling. Each of these factors necessarily constrains the free movement of the animal. Here we describe a promising new system for monitoring physiology and behavior in freely moving mice, with significant advantages in size, weight, cabling, signal quality, and customizability. The key advance is the use of ultra-miniature analog camera modules (180 mg, 5×5×5 mm), allowing for lightweight multi-camera headsets, finely-focused video signals, and single-ended signal connections for low-impact tethering.

We present two examples of different arrangements of multi-camera headsets for use with electrophysiology: a four-camera adjustable-focus headset integrated with an electrophysiology headstage, and an independent two-camera fixed-focus headset. We then discuss the pros and cons of different configurations and systems, and general guidelines for designing a specific implementation of this system. We also describe the detailed implementation of each arrangement, with additional step-by-step procedures included as appendices.

## Results

### Four-camera adjustable-focus headset

First, we describe a four-camera headset, designed as an attachment for the Intan 32-channel electrophysiology headstage. The four cameras provide an independent view of each of the two ears, an overhead view of the eyes, whiskers, and rhinarium, and a forward-facing view of the visual scene. Figure 1 and Supplemental Video 1 show an example of the signals from this four-camera headset as a mouse explores an object in an open-field arena. Pupil diameter, gaze direction, whisking, and rhinarial movements can be measured from the overhead camera, and ear movements can be measured from the ear-facing cameras.

**Figure 1.**
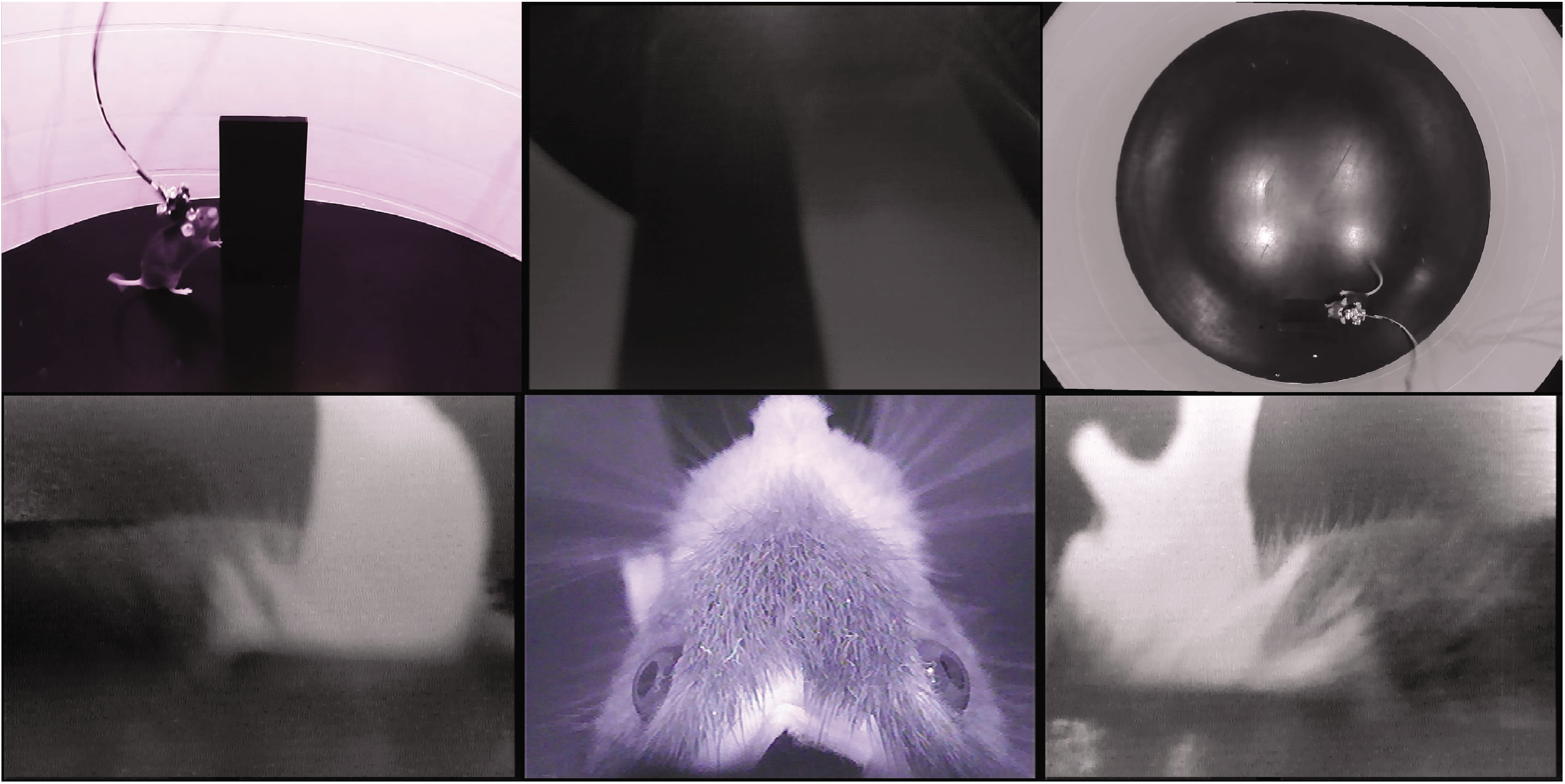
Example data from the four-camera headset. An example frame from Supplemental Video 1, showing the webcam views of a mouse exploring an open arena, from the side (top left) and above (top right), and the video signals from the four-camera headstage-integrated headset: left ear camera (bottom left), right ear camera (bottom right), overhead camera (bottom middle) and forward-facing camera (top middle). Note that the overhead camera video signal is of higher quality than the front-facing or ear-facing videos, due to their filtering.

Including the electrophysiology headstage, this camera headset weighs a total of 2.6g, and connects to implants weighing ~1.4g. Implants consisted of a Neuralynx electrode interface board (EIB), a 3D-printed base (Figure 2Ai), and a protective cap to allow for social housing (Figure 2Aii). Camera headsets consisted of analog camera modules assembled on a 3D-printed camera carrier (Figure 2Aiii), which attaches to the Intan headstage (Figure 2B,C).

**Figure 2.**
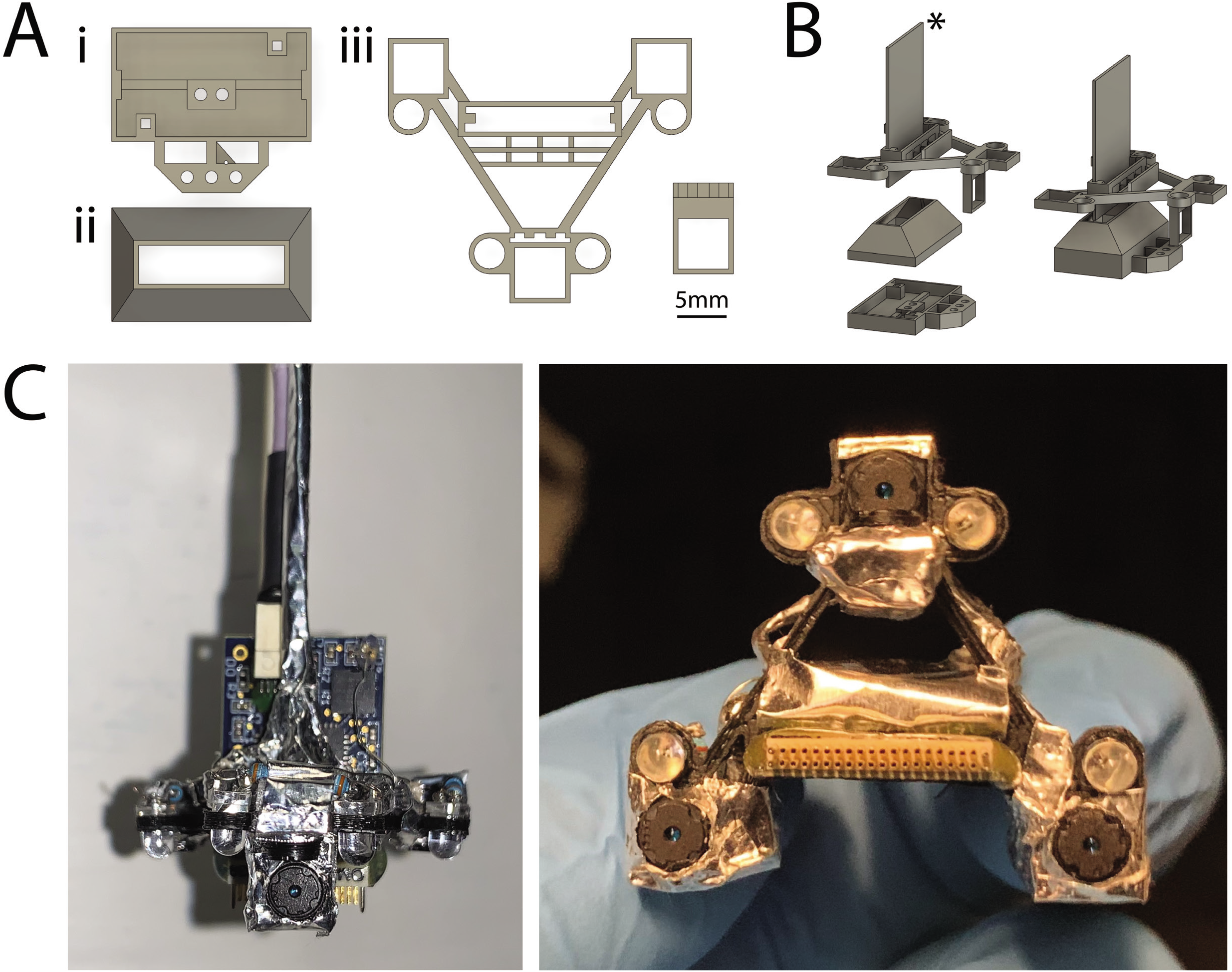
Four-camera headset design. A: 3D-printed components used for implants and headsets. i) Implant base that is implanted on the head. A Neuralynx EIB is attached to the implant base. ii) Protective cap that snaps onto the implant to allow for social housing. iii) Camera carrier that holds the cameras and their associated LEDs. The ear-facing cameras are each paired with a single LED for illumination, whereas the overhead camera has two LEDs (for illuminating the left and right side of the mouse’s face), and the forward-facing camera does not have an LED. The forward facing camera holder is attached to the main camera carrier component after printing. B: Rendered view of the components in A, separately at left and assembled at right. * indicates the RHD 32-channel electrophysiology headstage. C: A view of the headset assembled around the headstage from the front (left) and underneath (right).

Integrating cameras with the headstage in this way allows both electrophysiology and video signals to be collected with a single mechanical attachment to the head (at the EIB), and reduces the overall weight. This design also allows for stereotactic precision and reproducibility of camera views across animals and implants by combining the fixed dimensions of the implant with the stereotactic targeting of the electrodes. Additionally, the focal plane of each camera can be easily adjusted to accommodate for surgical or anatomical jitter encountered between animals (see “Reproducibility and precision” below for more details). When the factory-installed lens on the camera modules is maximally extended, the field of view is approximately 12 x 9 mm at the level of the focal plane, which is approximately 12 mm away. By screwing the lens all the way in, the camera’s focal length increases towards infinity, with a corresponding increase in field of view.

Each camera requires three electrical connections: a power, ground, and signal. With this headset design, the four cameras share power and ground connections, thereby only requiring a six-conductor tether (or six individual wires). The common ground introduces interference in the video signals caused by high-frequency crosstalk, but this interference can be removed through the use of simple low-pass filters (see “Powering and signal conditioning” below for more details). One camera can remain unfiltered, yielding a higher quality video signal. We chose the overhead camera in this case, to achieve finer detail of the eyes, whiskers, and rhinarium.

This headset design has some drawbacks compared to the two-camera headset described below. Because the headset is fully integrated, if one component were to fail, it may be difficult to individually replace it without rebuilding the entire headset (although fortunately the camera modules can be reused). Once assembled, the camera positioning is not adjustable, requiring iterations of building and testing to optimize camera angles for each use-case. Thought should be given to the experimental environment at this point in the design process as well. For example, the placement of the overhead camera in this design may be incompatible with some nosepoke designs, which would require redesign of either the headset or nosepoke. Lastly, while this headset arrangement allows for a large view of multimodal behavior, the overhead perspective may miss a portion of the pupil in rare cases of extreme ventral gaze.

### Two-camera fixed-focus headset

Next, we describe a two-camera headset designed to consistently capture the entirety of both pupils, and with higher detail. The headset consists of two bilateral cameras directed at the eyes, with a fixed focus achieved by the addition of a collimating lens (Figure 3A). To provide the lowest noise and highest-quality video signal, both cameras have independent signal, power, and ground connections (see “Powering and signal conditioning” for more details).

**Figure 3.**
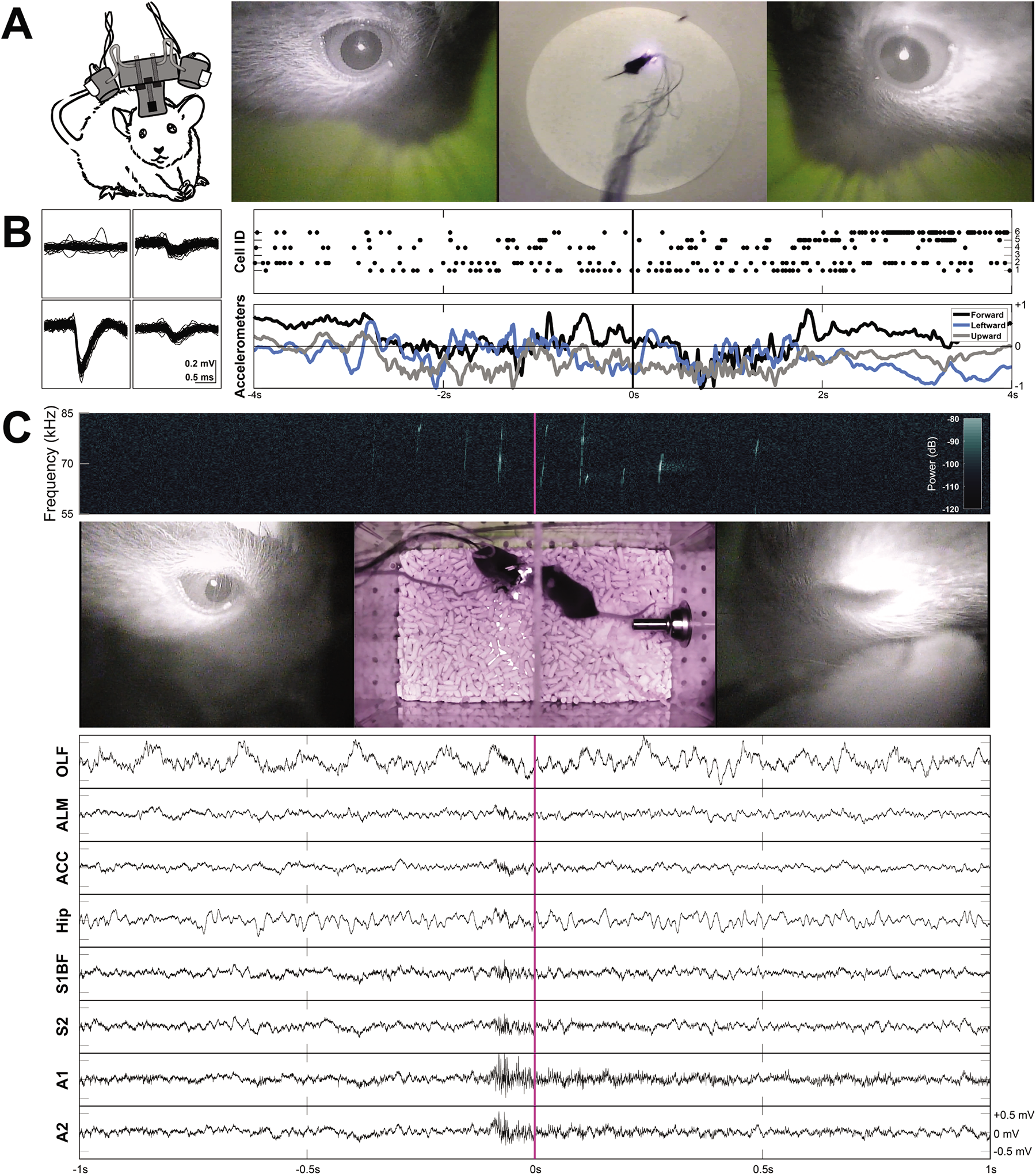
Two example applications of the two-camera setup with freely-moving behavior and electrophysiology. A: The two-camera fixed-focus headset on a mouse, without an electrophysiology headstage (left), and an example frame from Supplemental Video 2 showing the signals produced by this headset (right). B: Example waveforms across the four channels of a tetrode shown for an individual unit (cell ID 1) recorded during prey capture behavior. Rasters of cells firing and the accelerometer channels are plotted together (right) for the corresponding frames shown in A. C: An example frame from Supplemental Video 3. A spectrogram (top) shows ultrasonic vocalizations occuring during courtship behavior. Continuous LFP traces from various recording locations are shown below. OLF: olfactory bulb, ALM: anterior lateral motor cortex, ACC: anterior cingulate cortex, Hip: CA1 of the hippocampus, S1BF: barrel field of primary somatosensory cortex, S2: secondary somatosensory cortex, A1: primary auditory cortex, A2: secondary auditory cortex.

Although this headset is not integrated with an Intan headstage, it can be used with one. Example waveforms and rasters of unit activity recorded in anterior lateral motor cortex (ALM) during prey capture behavior are shown in Figure 3B and in Supplemental Video 2. Single neuron recordings were stable across many days. Raw continuous traces of local field potentials (LFPs) recorded from eight different brain areas during courtship behavior are shown in Figure 3C and in Supplemental Video 3 (see “Implants,” “Surgery,” and “Experimental conditions” for more details).

When focused on the surface of the sclera, these cameras can detect cyclotorsion of the mouse eye. Examples of this behavior can be seen in Figure 4 and Supplemental Video 4, which reveal torsional rotations of up to 11° in extent, at rotational rates of up to 4.6°/s. During periods of active movement, these rotational events occurred frequently, on average at 0.3 events per second.

**Figure 4.**
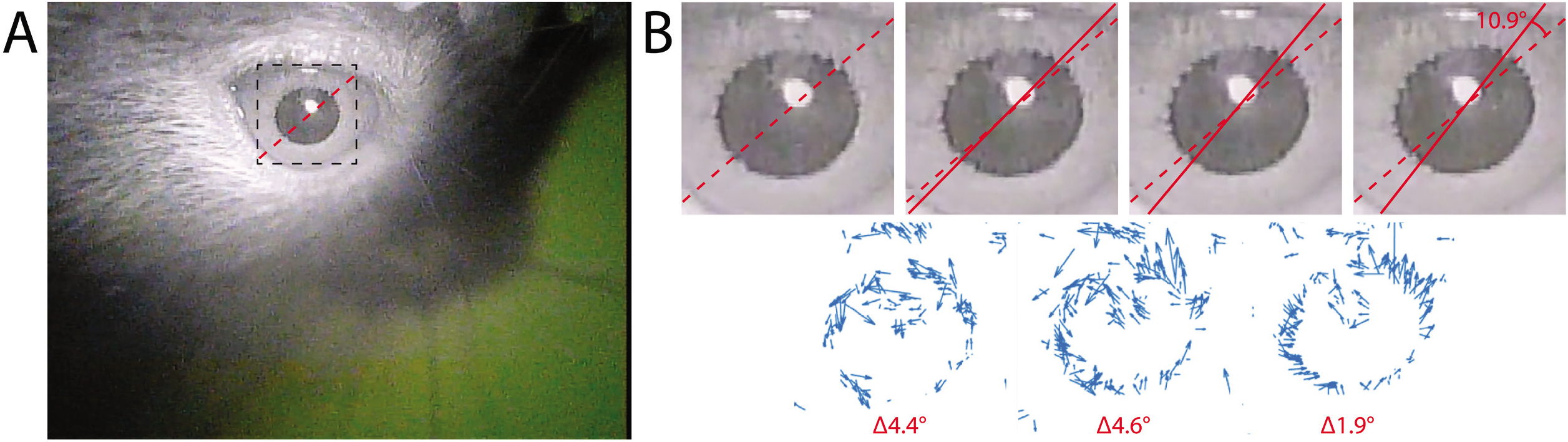
A: A deinterlaced frame from Supplemental Video 4, showing the right eye recorded with the two-camera headset. The inset marks the cropped location of the photos shown in B. B: Enlarged images of the pupil from four consecutive deinterlaced frames of Supplemental Video 4 at 60 Hz, with optic flow vector fields between successive frames shown underneath. Dashed red lines mark the orientation of prominent serrations across the pupillary rim of the iris in the first frame. Solid red lines track these serrations in consecutive frames. See Supplemental Video 4 for the full video. Optic flow was calculated in Matlab using the Lucas-Kanade derivative of Gaussian method with a noise threshold of 0.0005, and plotted with a scale factor of 100.

We tracked cyclotorsion using the unique pattern of natural serrations of the mouse eye in a manner similar to iris-registration software used to track torsional movements in humans during laser eye surgery (Shen et al., 2010). These serrations appear above the lens, at the pupillary rim of the iris, and vary between individual mice. Tracking these serrations allows torsional position to be determined without the use of more invasive techniques, such as limbal markings (Shen et al., 2010), scleral search coils (Robinson, 1963), or other magnetic implants (Schwarz et al., 2013).

Natural cyclotorsion of the eye has been observed in pigeons (Benjamins and Huizinga, 1929), chickens (Schwarz et al., 2013), and primates (Wells, 1794; Helmholtz, 1925), but to our knowledge has not previously been described in mice. Cyclotorsion is known to interact with the vestibular system to provide gaze stabilization as the head rotates about an axis of fixation or an object of smooth pursuit (Tweed and Vilis, 1987; Crawford et al., 1991; Angelaki and Dickman, 2003). The occurrence of cyclotorsion events up to 11° in mice highlights the importance of capturing all eye movements for studies of vision in freely moving animals (Ballard, 1991), and its necessity for studies that seek to relate retinotopic receptive fields to visual stimuli or behavior (Cooper and Pettigrew, 1979).

The mounting system of this headset uses steel wire and cement, rather than a 3-D printed carrier, which is easier for initial setup, camera angle adjustment, and troubleshooting, but lacks stereotactic precision, stability, and reproducibility across animals and across headsets. In other words, it’s a good choice for pilot experiments. Supplemental Video 5 illustrates how this mounting system can be used for testing different camera angles in pilot experiments (in this case, as a mouse performs a jump). The independent mounting system and additional lenses reduce the weight efficiency of this headset, but with only two cameras this is not a major concern. The camera angles and focal distance can be adjusted by bending the steel mounting wires, allowing customization to individual mice, but this adjustment method is not well-suited for repeated daily customization of a single headset to multiple different mice. Ultimately, once the spatial arrangements of the components have been optimized, the headset could be adapted to use a 3-D printed carrier, similar to the four-camera headset described above.

For either headset configuration, the headstage can provide power and ground connections of the correct voltage for the cameras, reducing the tether requirements even further, but at the cost of introducing appreciable levels of 60 Hz noise into the electrophysiology channels. The level of noise with headstage-provided power is unacceptable for LFP recordings, and for small-amplitude single neuron recordings, but could allow isolation of large-amplitude single neurons.

## Discussion

Animals move a lot. Whether these are active sensing movements such as saccades or whisking, or movements directed at another goal such as capturing prey or returning home, the effects on sensory input are profound. Movement also has a substantial impact on brain activity even beyond the directly-involved sensory and motor areas (Musall et al., 2018; Stringer et al., 2019; Salkoff et al., 2020). To understand how the brain underlies both sensation and action during freely-moving behavior, whether natural behavior or choice tasks, will therefore require precise measurement of the movements of the eyes, ears, and whiskers. Here we have described a head-mounted multi-camera system for recording simultaneous video of multiple body parts along with high-density electrophysiology in freely-moving mice. The core components are lightweight (180 mg) and inexpensive (US$23) analog camera modules. We show that this system provides high-precision tracking of eye movements, such as cyclotorsion, as well as movements of other body parts. This allows alignment of neuronal activity with motor events, including both active sensing and general body movements, in freely-moving animals.

### Size and weight

The size and weight of head-mounted devices is critical for unobstructed movement. At 5×5×5 mm and 180 mg, the size and weight of these camera modules serve to minimize their effects on mouse behavior. The four-camera headset described above, including the electrophysiology headstage, weighs a total of 2.6g, and connects to implants weighing ~1.4g, for minimal effects on natural behaviors such as prey capture and vocal interaction. Figure 5 shows the differences in size and weight between the LeeChatWin RS-306 camera (with and without an optional collimating lens) and the Adafruit 1937 camera, a leading alternative for behavioral monitoring of freely-moving mice (Meyer et al., 2018). While the RS-306 camera is significantly smaller and lighter, the Adafruit 1937 camera has superior temporal resolution, capable of capturing digital 640×480p frames at 90 Hz. The Adafruit 1937 can also be configured to capture 1080p frames at 30Hz, or 720p frames at 60Hz.

**Figure 5.**
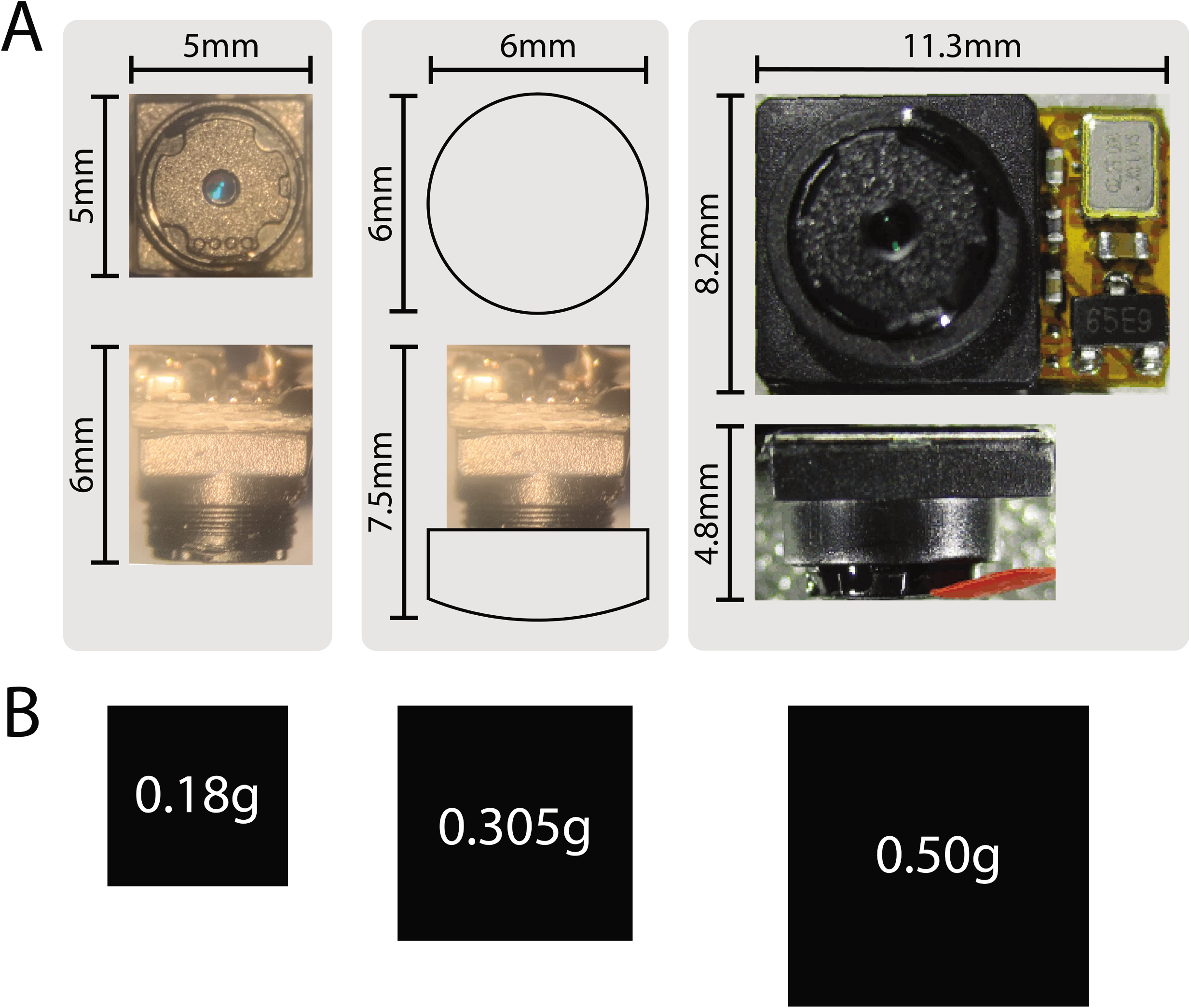
A: Size comparisons between the camera modules used in the four-camera headstage-integrated headset (left), two-camera headset with additional collimating lens (middle), and an Adafruit 1937 picam camera module (right). The factory-installed adjustable lens has 1 mm of travel, and is shown maximally extended (left, for a total camera depth of 6 mm), and minimally extended (middle, for a camera depth of 5 mm, or 7.5 mm with the additional collimating lens). Broader views of the camera modules are also shown in Appendix 1 and 2. B: Weight comparisons represented by area for the corresponding camera modules shown in panel A.

### Tethers

For multi-camera video, tethers will likely remain necessary until further advances in power and high-bandwidth telemetry or local data storage become available. The effect of the tethers will therefore remain a hindrance to freely-moving behavior of the mouse. As a mouse turns, the tether accumulates torque which strains their free movement. Importantly, since this system uses individual wires for signal and power connections, as opposed to the flat ribbon cables of the Adafruit 1937 camera, they can readily be integrated into slip ring commutators and pulleys to relieve strain from the tethers, which may be more difficult with a ribbon cable. Using modern commutators which utilize close-loop systems to provide active accommodation could then be implemented to relieve any strain of the electrophysiology and camera tethers, as well as optical fibers (Hoshino et al. 2020). Active counterbalancing of the tethers through closed-loop tracking and motorized pulleys should also be possible. Active systems might be necessary if passive slip ring commutators and pulleys present enough friction that mice are affected by accumulated torque before it is relieved by the device. We have not yet tested slip ring commutators or pulleys with this system.

### Reproducibility and precision

The precise positioning of electrodes and cameras is limited by surgical jitter of implants and individual anatomical differences across mice. The solution we described here is to integrate conventional stereotactic targeting with 3D-printed implant design. A promising framework for these types of implants is the RatHat, which is an open-source self-targeting 3D-printed brain implant system (Allen et al., 2020). Manual adjustment of the fine focus of the individual camera modules can also help compensate for remaining differences in focus encountered between animals. Taking note of the markings on the face of the lens when adjusting between animals is helpful for reproducibility in this case.

### General design guidelines

Other undesirable consequences of all head-mounted devices include partial occlusion of the visual field, interference with sound localization due to sound shadows, and interference with access to apparatus components such as nosepokes. For these reasons, camera headsets should ideally be designed with specific experiments in mind, to balance the benefits of the desired behavioral and physiological signals with the impact on sensation and behavior.

When customizing the camera set-up to target different camera views, electrode locations, or other hardware, we recommend the following design considerations. First, determine the locations of optical fibers or microinjection ports, which require open access during experiments. The EIB should then be placed where it doesn’t impede this access. We typically place the EIB directly over electrode arrays. A different choice of EIB, depending on the headstage used, may affect this placement. The implant base should then be designed and/or positioned to accommodate the EIB and other hardware. Finally, design the camera carrier based on how it will mount to the implant base and/or EIB, in order to target the cameras at the desired viewpoints.

### Future directions

We envision a number of possible extensions of this system. For example, by finely focusing on the sclera, we were able to measure large and frequent cyclotorsion events. In addition to azimuth and elevation, this provides the final degree of freedom required to reconstruct the visual scene in retinotopic coordinates. If combined with modern inertial measurement devices and video tracking, this feature would make it possible to retinally map the visual field of freely-moving mice in 3-dimensional space. We have not explored the upper limits on the number of camera modules that can fit on the head and be supported by a mouse without noticeably impacting behavior. Improvements in implant and tether design will improve efficiency and will likely increase the number of devices that can be included. The use of high-density silicon probes, such as neuropixel arrays (Juavinett et al., 2019), would dramatically increase the number of neurons that can be recorded alongside head-mounted video. Optical imaging of neural activity using these sensors may also be a promising approach. A widefield miniscope using this analog camera module could be integrated with behavioral camera arrays, like those in Figure 1, to investigate large-scale brain activity in freely moving conditions.

## Methods

### Preparation of camera modules

We used LeeChatWin RS-306 miniature analog camera modules (Part #1), but other miniature analog camera modules with similar specifications are available from a range of manufacturers. Before modifying the cameras in any way, we ensured that they produced a clean and stable video signal out of the box by powering the camera’s attached barrel cable with a 12V DC power supply and connecting its RCA composite video connector to a display. We then removed the infrared filter as described in Appendix 1, cleared away excess rubber as described in Appendix 2, and stripped insulation from the leads as described in Appendix 3.

### Four-camera adjustable-focus headset

#### Construction

All 3D-printed components (Figure 1) were designed with Fusion 360 software (Part #2), prepared using Cura (Part #3), and printed with polylactic acid (PLA) on a Monoprice Maker Select Plus 3D Printer (Part #4).

We 3D-printed a camera carrier (Part #5) as shown in Figure 1Aiii, and in Figure 6 with numbered labeling. The carrier was inspected and tested to ensure the camera modules would properly fit into each camera holder as described in Appendix 7. We then applied a thin coat of superglue to the grooves of the forward camera holder (marked 5 in Figure 6), and inserted it into the corresponding slot in the camera carrier (marked 6 in Figure 6, also see photo in Appendix 7).

**Figure 6.**
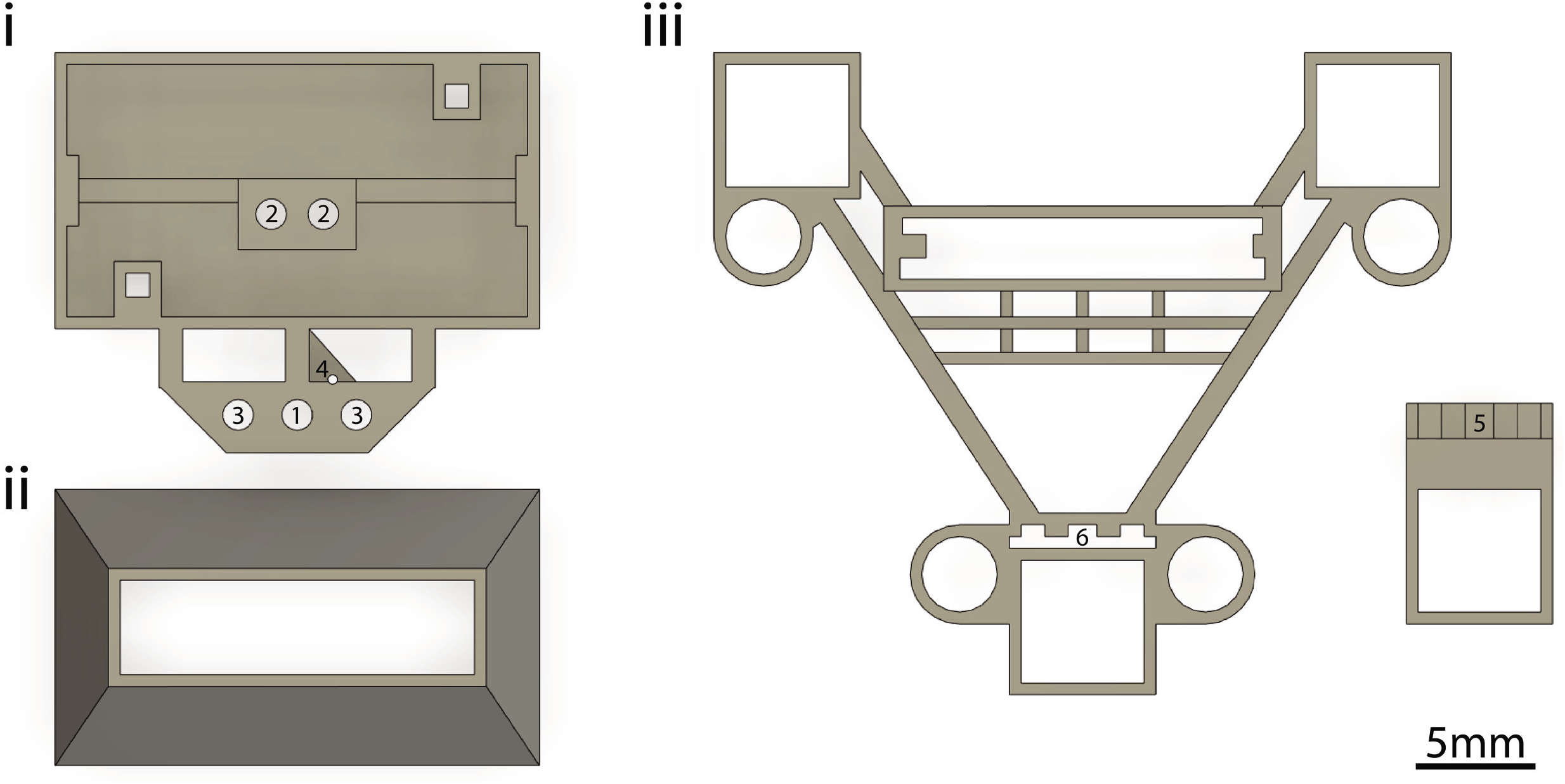
The 3D-printed components used for implants and headsets as shown in Figure 2, but shown here expanded and with numbered labeling. Numbers are referred to in the Methods (1-4: see Implants, 5-6: see Four-camera headset, Construction).

To create the headset, we first prepared a wiring harness with measured segments of wire needed to power the LEDs and camera modules on the headset (Appendix 4). We then soldered the prepared wires to each camera module (Appendix 5), shielded them with aluminum tape (Appendix 6), and cemented them onto the headset (Appendix 7). We then inserted LEDs into the receiver holes in the carrier and soldered the leads to their respective camera module (Appendix 8). The LEDs provide constant and sufficient illumination of the eyes, ears, or face during experiments regardless of head position. Next, we soldered the power wires to a single power tether, and the ground wires to a single ground tether (Appendix 9). We then applied the last piece of shielding (Appendix 10) and attached the headset to an Intan headstage by inserting the headstage into the receiver slot in the carrier, and then sliding the carrier all the way down onto the headstage (Figure 1B). A friction fit is sufficient to keep the headset firmly mounted to the headstage indefinitely. Repeated attaching and detaching of the headset to headstages could cause wear and tear and is not recommended. Finally, we electrically connected all the shields and pinned them to the local ground of the headstage (Appendix 12).

#### Powering and signal conditioning

We terminated the ends of the signal, power, and ground tethers with male jumpers for easy connection/disconnection. We used a +3.3V terminal on an RHD2000 USB interface board to power the cameras for use with or without accompanying electrophysiology, but any +3.3V power supply should work. The cameras could optionally be powered directly by the headstage as well, reducing the number of wires in the tether and still allowing accelerometer signals to be acquired, but at the cost of introducing noise on the electrophysiology channels.

The 4 cameras can share power and ground connections, thereby using only 6 wires, but the common ground introduces interference caused by high-frequency crosstalk. If additional wiring is not a consideration, this interference can be avoided by using independent power and ground connections for each camera (i.e. 12 wires for 4 cameras). Alternatively, the interference can be removed with simple low-pass filters. We constructed these filters on a separate breadboard for the camera signals, and integrated it into the system as shown in Figure 7. One camera signal can remain unfiltered, yielding a higher quality video signal.

**Figure 7.**
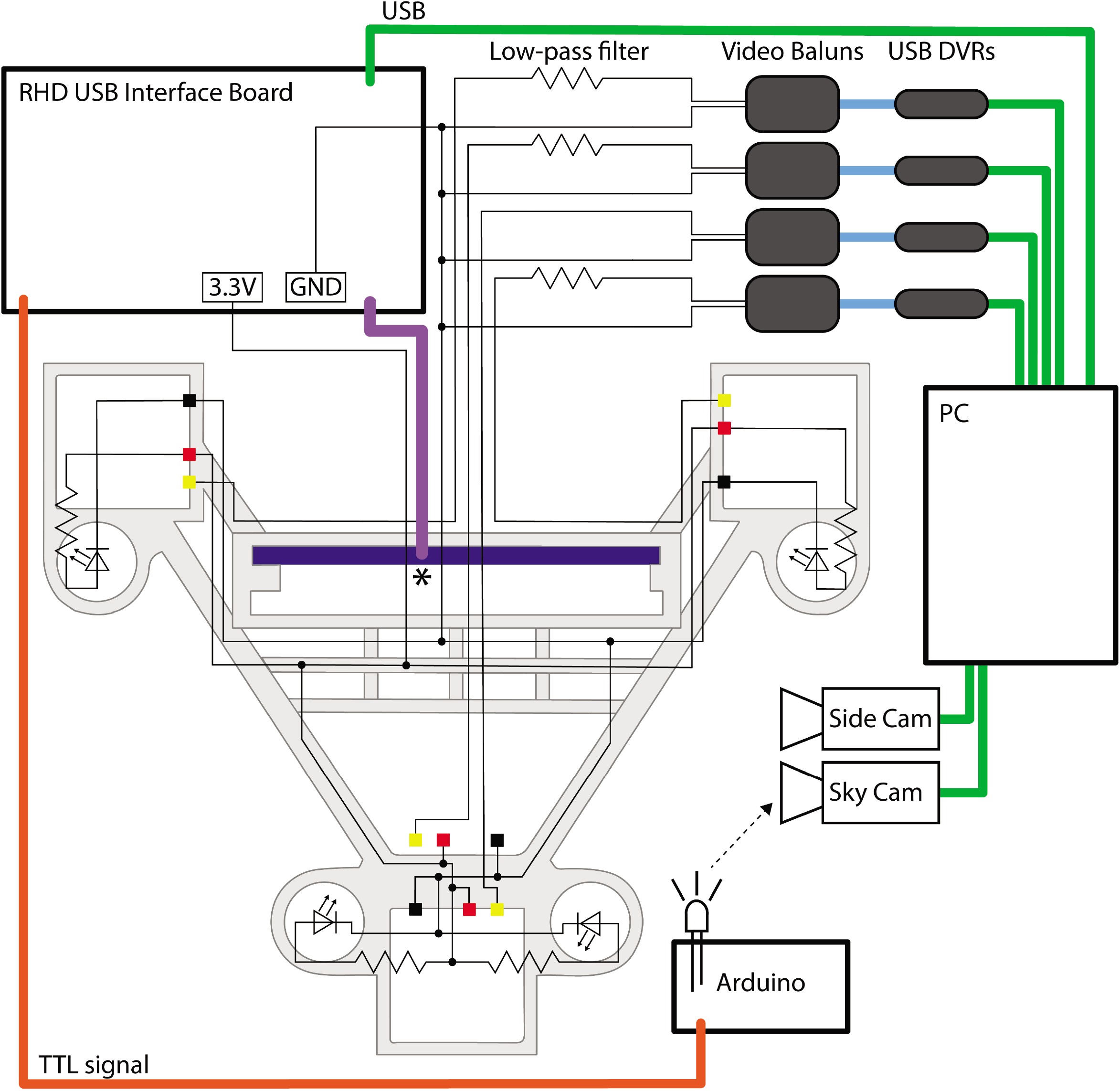
Circuit diagram and connections for the four-camera headstage-integrated headset. The camera carrier (same as Figure 1B) is shown in the background in grey. The RHD 32-channel headstage is represented by the dark blue rectangle, and is attached to its associated SPI cable represented by the purple line at the point marked *. TTL pulses sent to the RHD 2000 USB Interface Board are relayed through a BNC cable (orange) to an Arduino Uno for synchronization of electrophysiology and video signals by illumination of an IR LED in view of the webcam placed above the arena (the SkyCam). Yellow, red, and black squares represent the soldering locations on the camera modules for the signal, power, and ground wires. Low-pass filters are constructed on a breadboard, and are then connected to video baluns through a BNC connection. From the video baluns, the signal passes through a BNC to RCA connection (blue) to the USB DVRs. Green lines represent USB cables. Black dots indicate logical connections of signals and components; for true solder locations, see the instructions in the appendices.

The filtering eliminates the crosstalk between the cameras produced by the common ground loop, but comes at the cost of losing the chrominance signal and the high frequency portion of the luminance signal. Therefore, color information is lost, a reduction in the luminance of the signal causes a dimming of the video, and some minor noise artifacts such as dot crawl are introduced into the filtered video signals. The drop in luminance can be compensated for by increasing the intensity of LED illumination (by changing the value of the current-limiting resistor for each LED to produce sufficient brightness).

Finally, we pass the filtered signal and ground for each camera through individual video baluns (video signal isolators, Part #6) before being connected to USB video capture cards (DVRs) for digitization of the video signal. The video baluns were used to isolate a second ground loop that would otherwise normally occur at this point in this system, produced by having two paths to ground: one to the ground of the camera power supply and one to the ground of the computer.

### Two-camera fixed-focus headset

#### Construction

We first applied a small amount of super glue (Part #7) for strain relief between the wires at the base of the prepared camera modules so they were secured for long-term use. We then soldered an independent signal, power, and ground tether (Part #8) to each camera module. A 560 **Ω** resistor (Part #9) was soldered at this location to the power wire, as well as an additional wire to the ground tether, for powering an illumination LED (see below). Shrink tubing was then applied to these three junctions (Part #10). We then attached a 6 mm collimating lens (Part #11) onto the front of each camera module using tape (Part #12), wrapped it with a layer of micropore surgical tape (Part #13), and applied a thin coat of super glue.

We then prepared two 15G hypodermic tubes to a length of 5 mm, and a 20G stainless steel wire (Part #14) to a length of 2cm. The hypodermic tubes were inserted onto the front two screws of an assembled implant (3 in Figure 6), and the stainless steel wire was cemented between them with super glue, followed by a layer of dental acrylic. Once the acrylic cured, we then cemented the two camera modules bilaterally to the stainless steel wires. We then cemented 3 mm IR LEDs (Part #15) to the sides of each camera module in a coaxial fashion to provide epi-illumination of the eye, and connected the leads to the power (through the current-limiting resistor) and ground tethers. Supplemental Video 5 shows an example of the video signal that is produced if epi-illumination from an attached IR LED is not utilized.

#### Powering and signal conditioning

The distal ends of the camera signal, power, and ground tethers were then soldered back to their original connections on the factory-provided wiring harness with the power and video connectors. The two cameras were then each powered with an independent 12V DC power supply and connected with RCA cables to USB video capture cards (DVRs, Part #16) for digitization of the video signal. The factory-provided power connector includes a step-down from 12V to 3.3V.

### Data acquisition and processing

Electrophysiology and accelerometer signals were acquired from an attached headstage amplifier (Part #17) with an RHD2000 USB interface board (Part #18) using OpenEphys software (Part #19). We identified single neurons offline using Kilosort spike sorting software (Part #20).

For either headset design, DVRs were connected to a recording PC, and video signals were acquired with Bonsai software (Part #21) at 30 Hz and a resolution of 1280×960. Video from additional webcams (Parts #22,23) were also simultaneously acquired with Bonsai at 30 Hz and a resolution of 1280×960 or 1920×1080. We increased the brightness of the ear-facing and overhead camera videos using the color balance function in Bonsai. Video can also be flipped or rotated at this point; for example, we rotated the signal from the forward camera video to compensate for the installation orientation. We also used Bonsai to record timestamps for all captured frames and log them to individual csv files for each camera. To synchronize the video signals with electrophysiology and accelerometer signals, we positioned an IR LED in view of the webcam, and drove it with TTL pulses also recorded by the RHD2000 USB interface board. We note that the DVRs are susceptible to electrical interference, e.g. from nearby power supplies or equipment that can introduce artifacts into the digitized video signal if too close to the DVRs.

The RS-306 camera produces an NTSC analog waveform encoding 4:3 aspect ratio interlaced frames with an optical resolution of 800×480 at 30 Hz. We deinterlaced the video offline to remove combing artifacts and recover the full 60 Hz field rate. Analog video is interlaced such that each 30 Hz video frame consists of two 60 Hz fields taken in sequence: the first containing the odd lines of the image, and the second containing the even lines. We used a line-doubling deinterlacing algorithm to separate each frame into two consecutive deinterlaced images consisting of the odd or even field lines, and doubled line width to preserve image dimensions, yielding deinterlaced video at 60 Hz, with an effective single-frame optical resolution around 800×240. The horizontal and vertical angles of view are 60° and 45°.

Stereo audio recordings for vocalization experiments were obtained with two Brüel and Kjær 1/4-inch microphones (Part #24) and acquired with a Lynx 22 sound card (Part #25) and Audacity software (Part #26). The transducers of the microphones were positioned at the ends of the cage and angled at 45 degrees to point at the base of the interaction barrier. We delivered white noise bursts through a free-field speaker at the beginning and end of each experiment along with a TTL pulse to an IR LED and the RHD 2000 interface board to synchronize audio and video recordings with electrophysiology and accelerometer signals. For the spectrograms in Figure 3b, we included noise reduction processing in Audacity using a noise profile of 5 seconds, noise reduction of 19 dB, sensitivity of 24, and frequency smoothing of 12.

### Implants

We tapped the three holes at the front of the implant base (Figure 1Ai, Part #27) with a 00-80 tap, and cut a 00-80 screw (Part #28) to an 8 mm length and screwed it into the center hole (marked 1 in Figure 6) with a washer (Part #29) to serve as a drive screw. We then tapped a 6 mm piece of plastic to serve as a cuff, and mounted it onto the end of the drive screw to create a total height of 1.3cm from the bottom of the cuff to the top of the drive screw. For tetrode implants, we then inserted two 6 mm 18G hypodermic tubes into the center of the implant base (marked 2 in Figure 6) to serve as guide rails for vertical travel of the base when advancing the electrodes. Two additional 8 mm 00-80 screws were screwed in the remaining holes at the front of the implant base (marked 3 in Figure 6) from the bottom up, to serve as a site for attaching the camera headset if an independent camera attachment site was desired.

For tetrode implants, we inserted two 6 mm 29G hypodermic tubes through the hole in the base (marked 4 in Figure 6), and cemented them in place with dental acrylic. For stainless-steel wire arrays, we inserted 7 mm 19G hypodermic tubes and cemented them in place with dental acrylic near their relative stereotactic locations on the implant base to accommodate two teflon-coated stainless steel wires each (Part #30).

We used small EIB pins (Part #31) to electrically connect individual stainless steel wires to the ground and reference channels on a Neuralynx EIB (Part #32). An EIB (electrode interface board) is a miniature break-out board with a headstage connector (such as an Omnetics connector) and connection points for implanted microwires, to provide a robust interface between a headstage and electrode or tetrode channels. The remaining channels on the EIB were similarly connected to prepared tetrodes (Part #33) or stainless steel wires for tetrode and wire-array implants, respectively. Once connected, we coated these sites with silicone sealant (Part #34). The reference and ground wires were either routed through the drive posts for tetrode implants, or two 19G hypodermic tubes in the case of wire-array implants. The tetrodes or remaining stainless-steel wires were then routed through their respective hypodermic tubes, and the EIB was lowered and flushly screwed into place on the surface of the implant base with two screws (Part #35). Stainless steel wires routed through the same tubes were cut to slightly different lengths so that we could properly identify them during implantation and note their respective channels on the EIB. The implants were then sterilized in 70% EtOH before implantation. Protective caps (Figure 1Aii, Part #36) were attached to implants after surgery.

### Surgery

All procedures were performed in accordance with National Institutes of Health guidelines, as approved by the University of Oregon Institutional Animal Care and Use Committee.

For the experiments described here, we used C57bl/6 mice (n=8) and made craniotomies as described below; these can be customized to the specific needs of different experiments. Mice were anesthetized with isoflurane (1-2%). A craniotomy was created over the right hemisphere, where a skull screw (Part #37) was fastened into place and cemented with dental acrylic. Two additional craniotomies were then created at −1.5 AP, −2 ML, 0 DV and −2 AP, −2 ML, 0 DV for the ground and reference wires. For tetrode implants, a small craniotomy was created dorsal to anterior lateral motor cortex (ALM) at 2.5 AP, 1.5 ML, 0 DV (relative to bregma). For wire-array implants, craniotomies were similarly created for recordings in the olfactory bulb at 4.5 AP, 0.8 ML, 0 DV, ALM at 2.5 AP, 1.5 ML, 0 DV, anterior cingulate cortex (ACC) at 1.98 AP, 0.35 ML, −1.8 DV, CA1 of the hippocampus at −2.06 AP, 1.5 ML, −1.5 DV, the barrel field of primary somatosensory cortex (S1BF) at −1 AP, 3.5 ML, 0 DV, secondary somatosensory cortex (S2) at −1.5 AP, 4.5 ML, 0 DV, primary auditory cortex (A1) at −2.9 AP, 4.5 ML, 0 DV, and secondary auditory cortex (A2) at −2.5 AP, 4.2 ML, 0 DV. Sterile saline was immediately applied and maintained at these locations to prevent drying of the dura.

Proper positioning of the cameras, electrodes, and other hardware such as optical fibers or microinjection ports will depend on the body parts and brain areas being targeted. For the experiments described here, EIBs were stereotactically placed with the center of the surface of their base at −5.5 AP, 0 ML, 10 DV from bregma. This was achieved with the fixed dimensions of the implant by positioning the drive screw at 4 AP, 0 ML, 0 DV relative to bregma, and ensuring the total height of the implant was 1.3 cm from the bottom of the cuff to the top of the drive screw. This positions the EIB centrally over the head and the camera carrier such that the cameras are over the ears and the front of the head.

For tetrode implants, the tetrodes were trimmed at this point so they would pass just beyond the level of the dura when the implant was lowered into its final position. Once at the proper length, we applied a thin coating of antibacterial ointment to the tetrodes, made a small incision in the dura, and slowly lowered the implant and tetrodes into place. We then coated the remaining exposed portion of the tetrodes in vaseline, and cemented the vaseline, cuff, and surrounding surface of the skull with dental acrylic. We then positioned the ground and reference wires just below the dura in their respective craniotomies, and cemented them with dental acrylic along with the drive posts so that no portion of the wires were left exposed.

For wire-array implants, the stainless steel wires were splayed out laterally so the implant could be lowered into its proper position. We then raised the implant 1.5 mm dorsally, and positioned the hippocampal wire vertically just above the dura, and cemented it to the hypodermic tube in this position with dental acrylic. This process was then repeated for the remaining electrodes targeting non-superficial structures. We then made small incisions in the dura at these locations, and slowly lowered the implant into position. We applied dental acrylic to these two sites and the cuff to cement them in place. Once the acrylic sufficiently stabilized the implant, the remaining wires were placed just below the dura in their respective craniotomies, and cemented with dental acrylic so that no portion of the wires were left exposed.

We administered Ketoprofen (5mg/kg) to reduce postoperative inflammation. The bottom surface of a protective cap (Part #36) was then coated with silicone sealant (Part #34) and snapped over the EIBs of the implant to cure in place. Mice were then given 7 days of postoperative recovery.

### Experimental conditions

Vocalization experiments occurred in the home cage of individually housed male mice, placed in an electrically shielded sound-attenuating chamber. A webcam (Part #22) recorded video directly above the arena (referred to as the “SkyCam”), with illumination from an IR LED light source (Part #38) and 6000K LED Floodlight (Part #39). We placed a perforated clear acrylic barrier (Part #40) between the resident male, who was unimplanted, and an implanted female. Vocalizations were recorded with two overhead Brüel and Kjær 1/4-inch microphones (Part #24).

Prey capture experiments occurred in a 24 inch diameter circular arena with a clean paper floor within an electrically shielded sound-attenuating chamber. SkyCam video of the arena was recorded as described above. Illumination was provided by an IR LED light source (Part #38) and 5500K lightbulb (Part #41). An implanted mouse waited in the arena until a cricket was remotely dropped into the arena, and then the mouse chased and captured the cricket (Hoy et al., 2016).

Free field exploration experiments occurred in the same arena as prey capture experiments. We used a webcam (Part #22) to record from one side of the arena, and a dome camera (Part #23) to record from above, with illumination provided by an IR LED light source (Part #42) and 6000K LED Floodlight (Part #39).

## Supporting information

Supplemental Video 3 (NormalSpeed)

Supplemental Video 3 (QuarterSpeed)

Supplemental Video 4 (NormalSpeed)

Supplemental Video 4 (QuarterSpeed)

## Acknowledgments

We thank Aldis Weible for help with implant design and attachment mechanics of the headstage-integrated headset; Kip Keller for help troubleshooting video ground loops; Iryna Yavorska and Jonny Saunders for technical assistance; and the Jaramillo lab for the use of their 3D printer. This work was supported by the National Institutes of Health, National Institute on Deafness and Other Communication Disorders Grant R01 DC-015828.

## Video Captions

Supplemental Video 1

An example video of the four-camera headstage-integrated headset showing the dynamic activity of the rhinarium, whiskers, eyes, and ears of a freely moving mouse. Panels show the webcam views of a mouse from the side (top left) and above (top right), and the video signals from the four-camera headstage-integrated headset: left ear camera (bottom left), right ear camera (bottom right), overhead camera (bottom middle) and forward-facing camera (top middle). Note that the overhead camera video signal is of higher quality than the front-facing or ear-facing videos, due to their filtering.

Supplemental Video 2

An example video showing rasters of unit activity from six cells in anterior lateral motor cortex (ALM) (bottom) during prey capture behavior (a mouse chasing and capturing a cricket). Traces from the accelerometer channels are shown below. Spikes from cells 1, 2, 4, and 6 are plotted in real time (waveforms at top) and indicated by 50ms tones of 800, 1200, 1400, and 1600Hz respectively.

Supplemental Video 3

An example video of courtship behavior. A spectrogram (top) shows ultrasonic vocalizations as they occur during the video of the behavior shown below. Continuous LFP traces from 8 brain regions are shown below. Audio was pitch shifted in audacity to bring the vocalizations within the frequency range of human hearing. Both a normal speed version of the video and a quarter-speed excerpt are shown. Note that this spatial arrangement of the cameras leaves ample room for mice to utilize the nosepoke ports of the interaction barrier.

Supplemental Video 4

An example video showing several epochs of natural cyclotorsion of the eye as a mouse moves around an environment. Both normal speed and quarter-speed versions of the video are shown.

Supplemental Video 5

An example video showing the pupil and whiskers during the natural jumping behavior of a freely-moving mouse. Using the mounting system described for the two-camera headset, different positions can be tested for pilot experiments, without having to iteratively rebuild headsets (compare the different camera angles in Supplemental Video 4 and Supplemental Video 5). This video also shows an example of the video signal obtained without an integrated IR LED (note how the illumination varies as the mouse moves relative to the overhead lighting).

## Appendices

## Appendix 0: Parts List

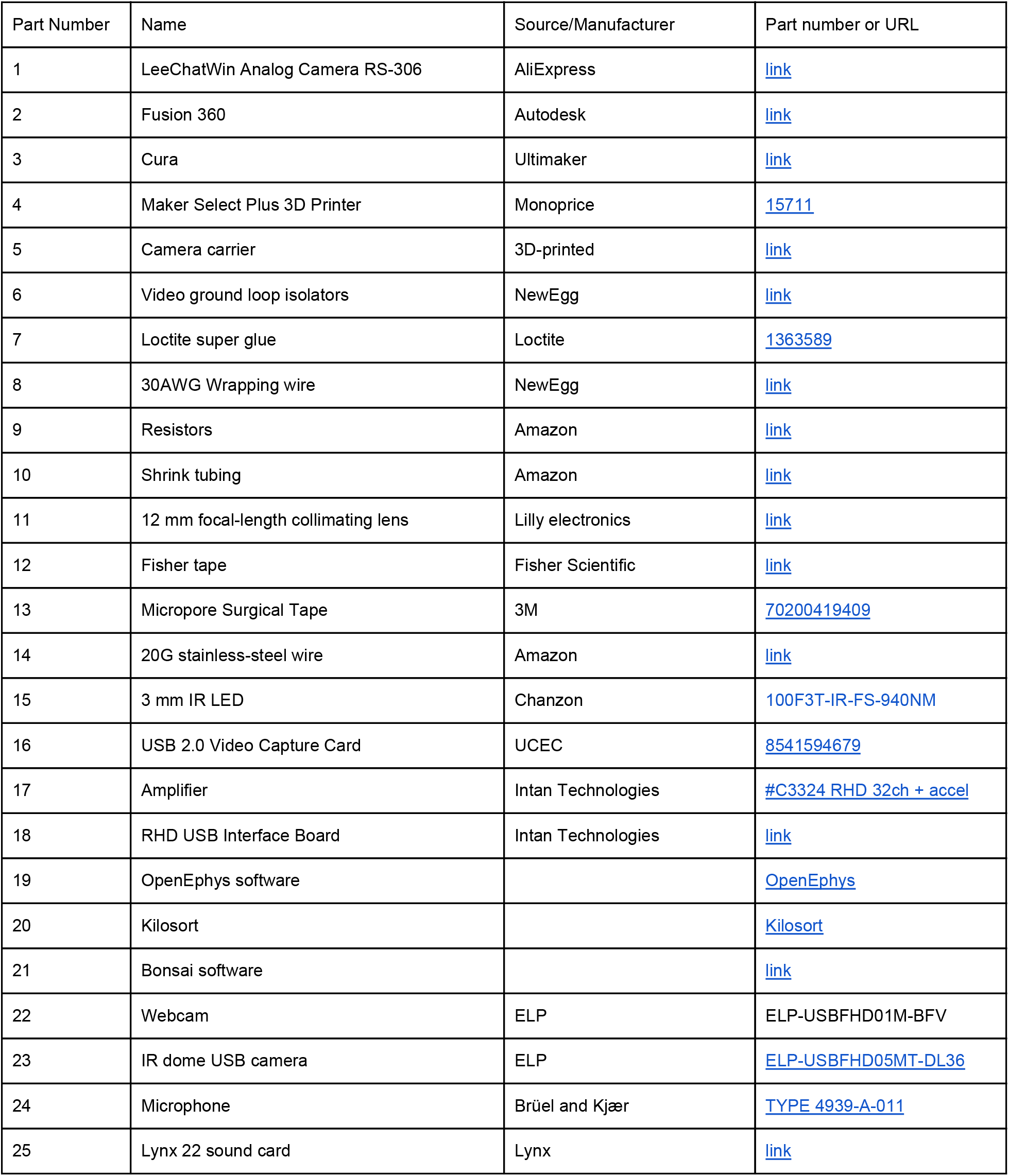
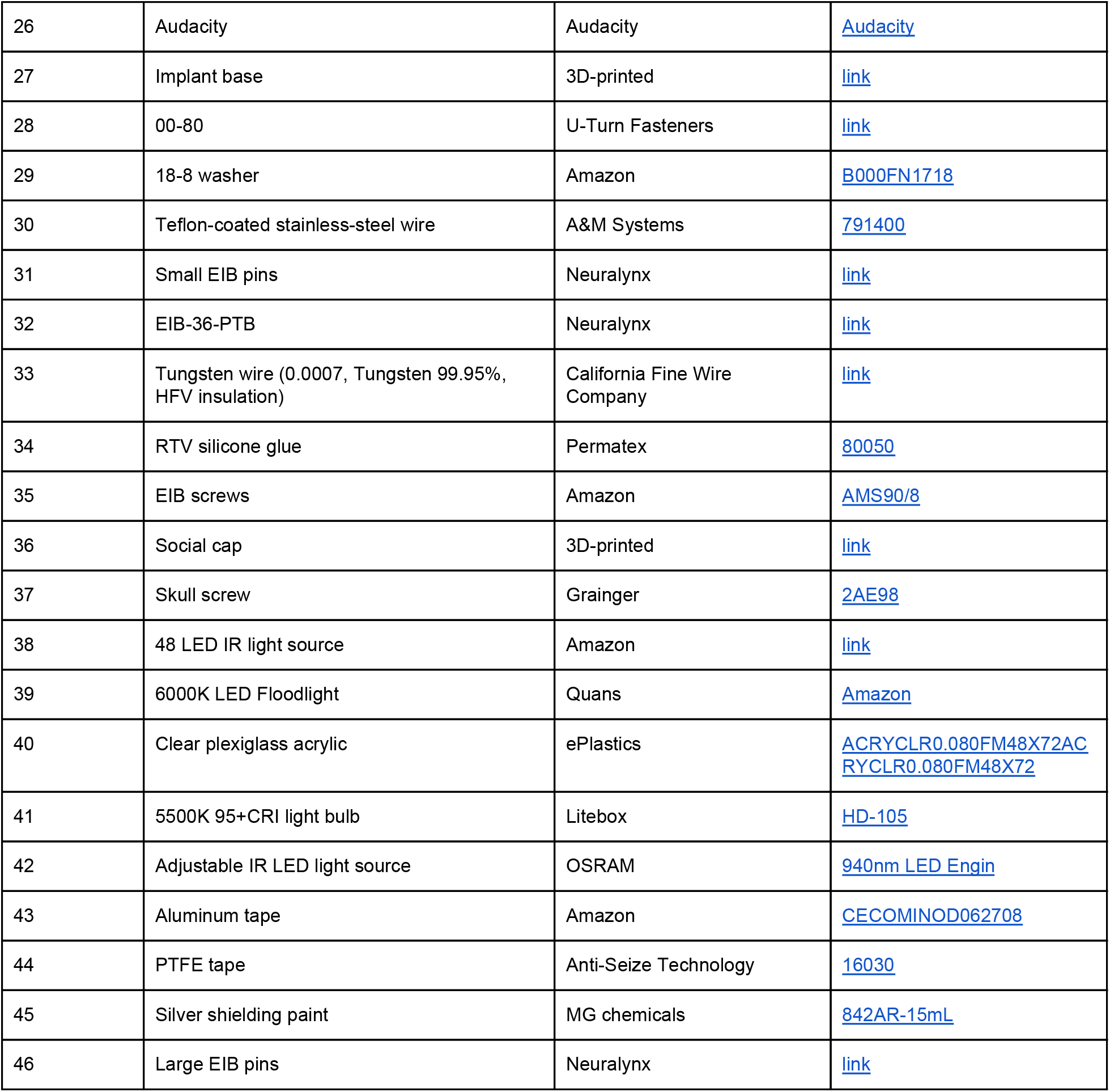

## Appendix 1: Removal of IR filter

Begin by unscrewing the lens located on the front of the camera (A). With the lens removed (B), place the camera out of the way and facedown on a clean surface to avoid any unwanted particles from accumulating on the sensor. Flip the lens over and begin using a scalpel blade around the edges of the filter to gently separate it from the lens. Once the filter has been sufficiently loosened around the edges, the blade will slide underneath it (C). Take care to avoid scratching the lens underneath during this process. Continue to work the blade around all sides of the filter until it can be fully removed from the lens (D). With the filter removed, screw the lens back onto the camera.

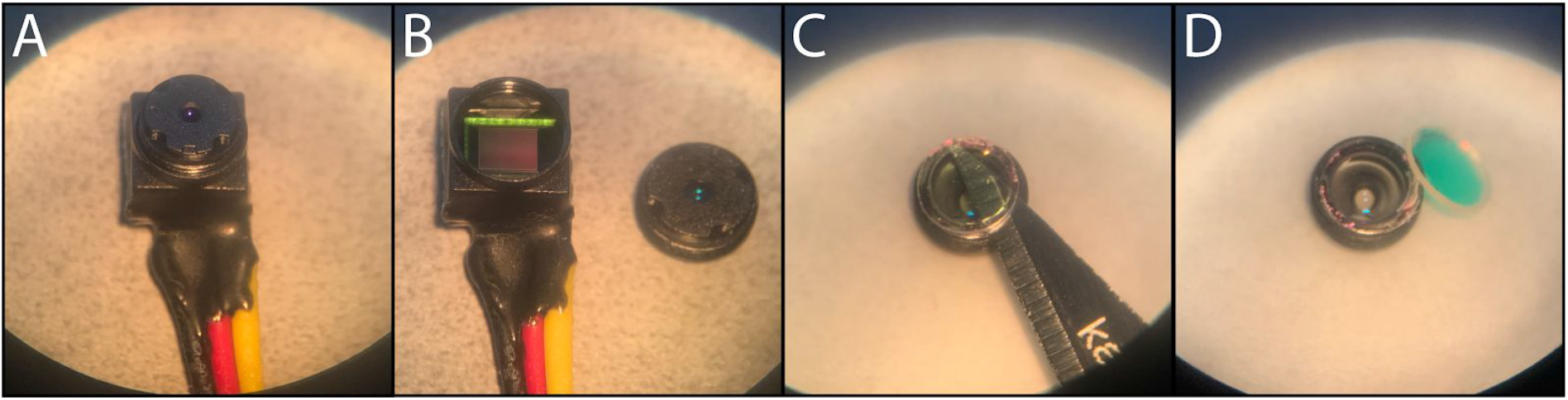

Test the video signal again using the camera’s power and signal connectors, to confirm that the signal is now sensitive to an infrared light source and that the lens and sensor are free of any debris or damage that affects the video signal.

## Appendix 2: Clearing excess rubber adhesive

The cameras often come with a small amount of external rubber adhesive (an example is shown in A), which can interfere with positioning when placed into a camera carrier. To prevent irregular placement, we recommend that you ensure that the four sides of the camera are clear of any rubber adhesive around the square-shaped core of the module. Gently remove adhesive using a scalpel and forceps until all sides of the camera are clear (as in B).

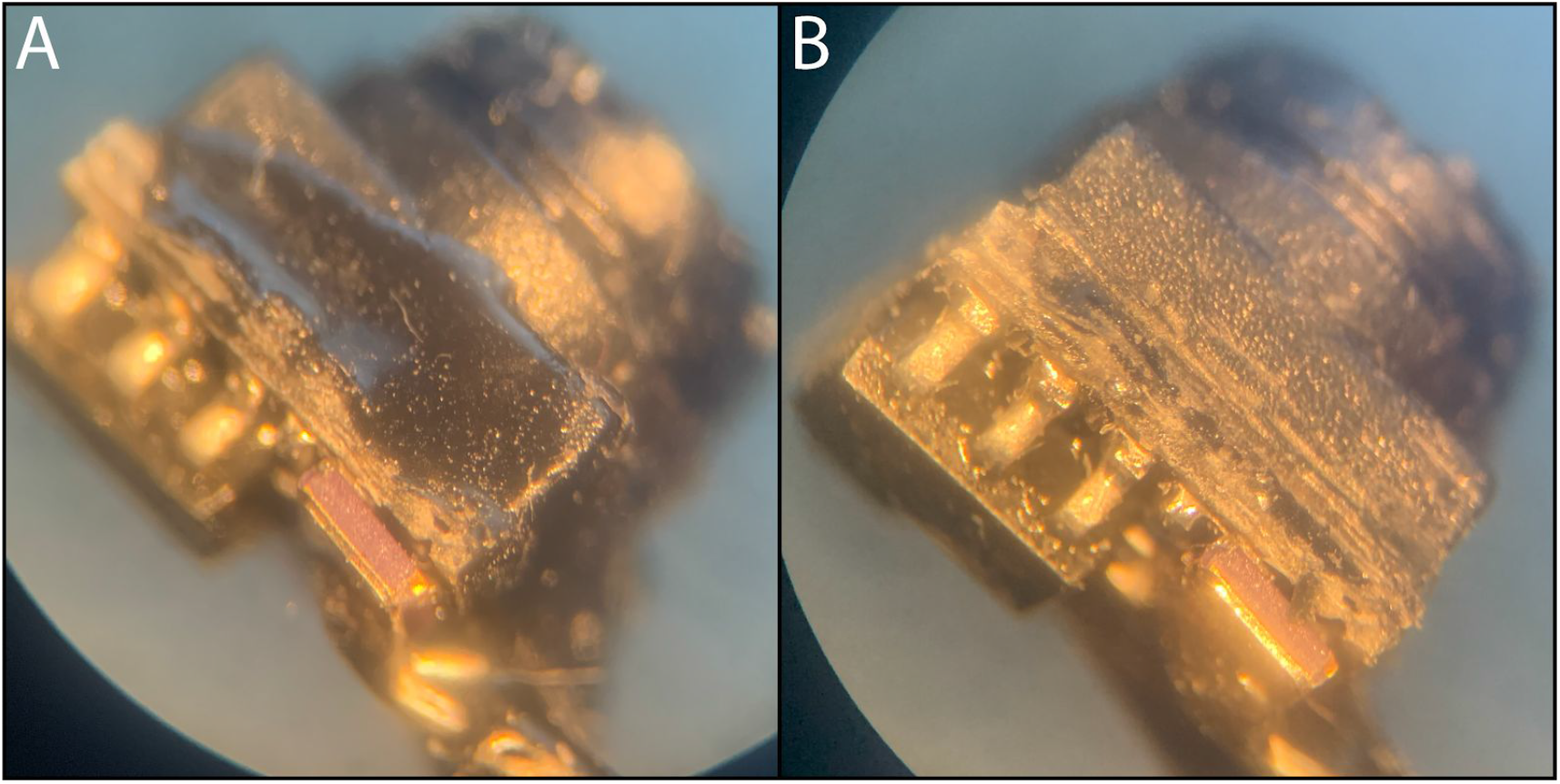

## Appendix 3: Stripping the camera module leads

Use wire cutters to cut the camera module’s signal, power, and ground wires at least 5 mm from the camera modules. Then, using a scalpel blade and forceps, carefully strip the signal, power, and ground wires to the base of the camera module (B). Take care when stripping near the base of the module, and leave some excess rubber around their connections, so they are less likely to break from stress during assembly.

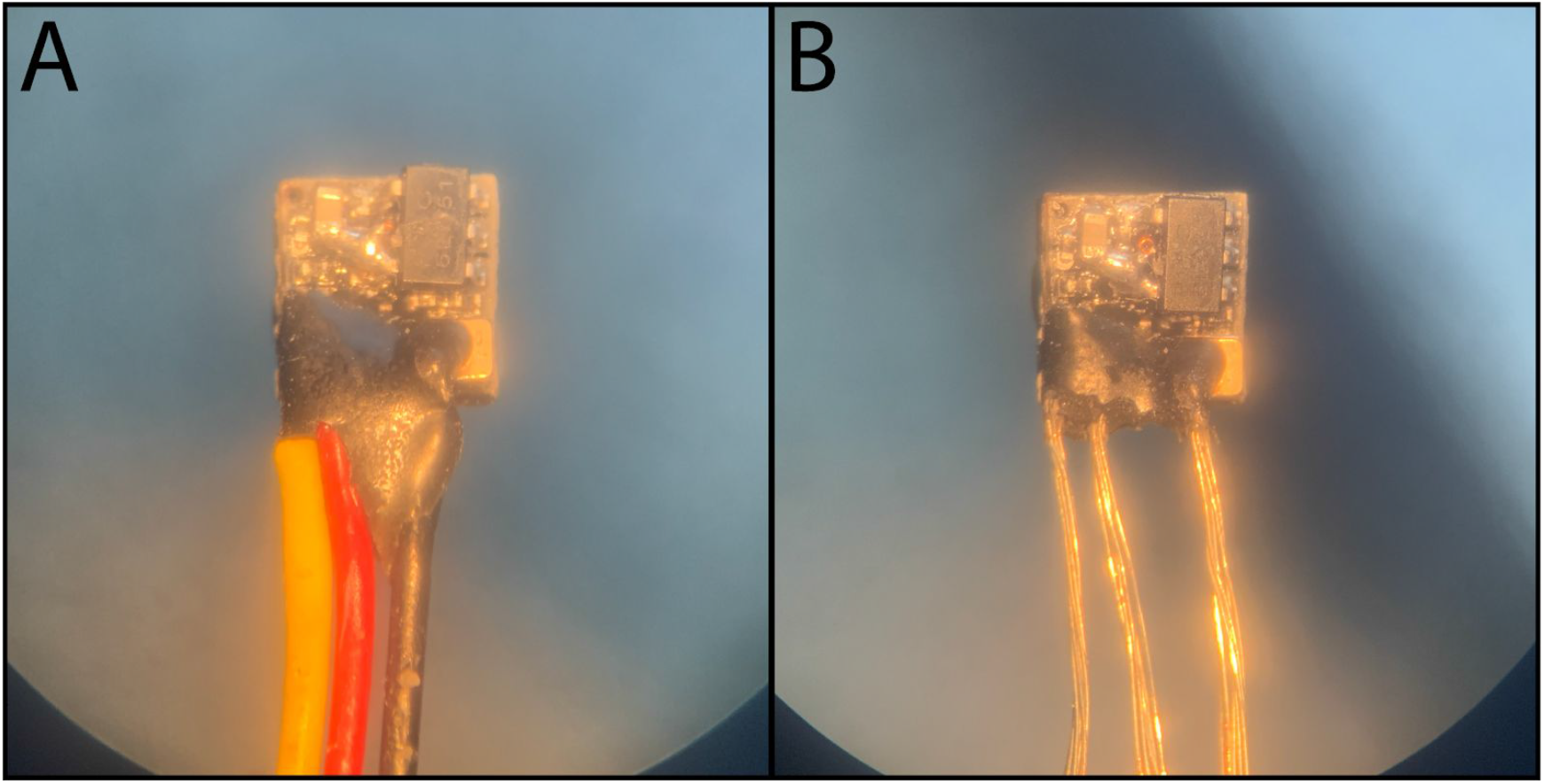

## Appendix 4: Preparing wire segments for soldering onto the cameras

Place the cameras in their appropriate holder on a carrier, with the accompanying LED(s) in place to use for reference while you prepare the wires that will power them. Strip and cut wires to the appropriate lengths, and solder on resistors where necessary (e.g. 390 **Ω**, Part #9). Panel A shows an example of power and ground wires customized for a four-camera headset, with specific gaps in insulation to solder to the cameras. Panel B shows an enlarged view of the power and ground wires for a right-ear camera. The signal wires for each camera (not pictured here) should be cut to be as long as your desired tether length.

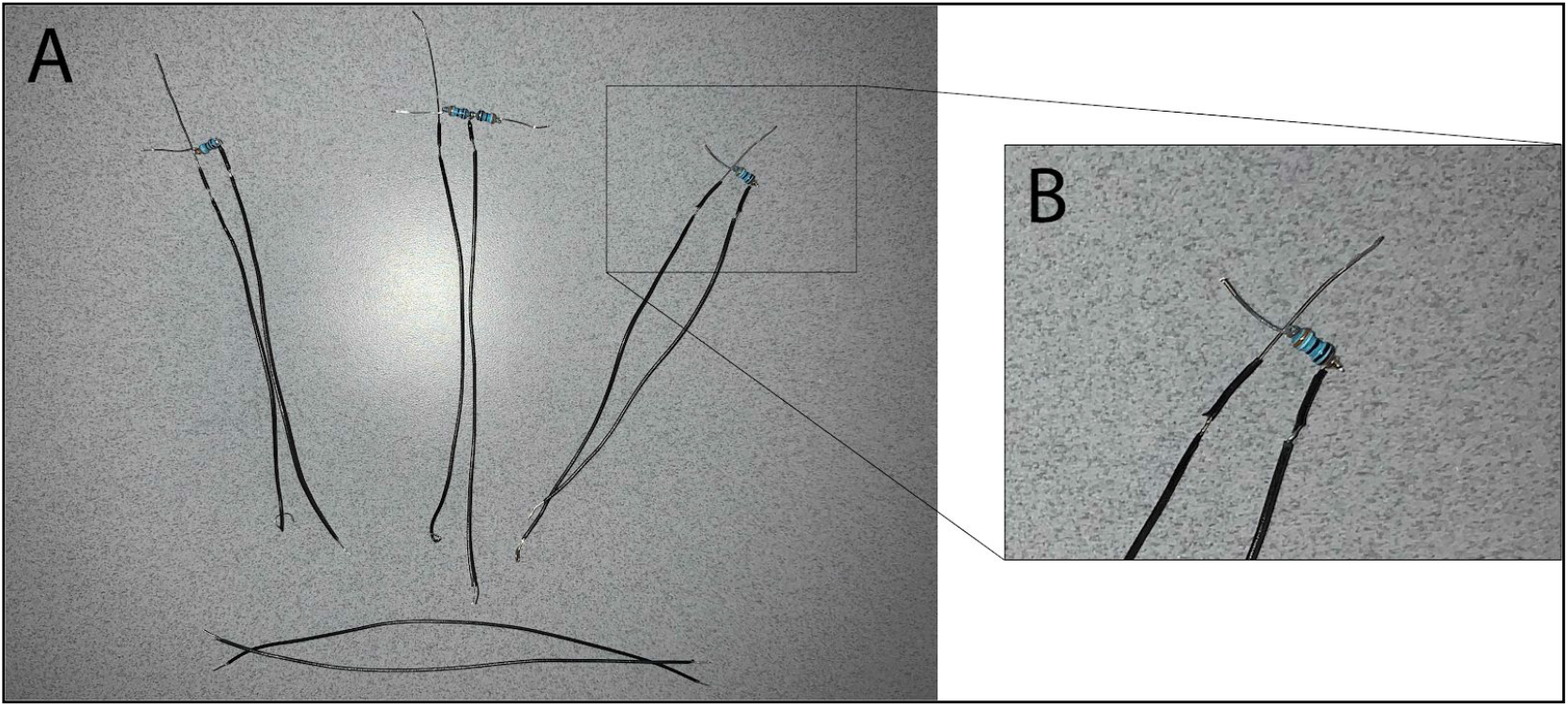

## Appendix 5: Soldering the wires onto the cameras and insulating them with silicone glue

With a camera in its proper position on a carrier, solder the signal, power, and ground wires, so they are oriented in the proper direction towards the center of the carrier. This is shown for a left ear camera as an example in panels A, B, and C. Once soldered and trimmed, remove the camera from the carrier and coat the exposed wire with silicone glue (D) and allow it to dry, so they are fully insulated.

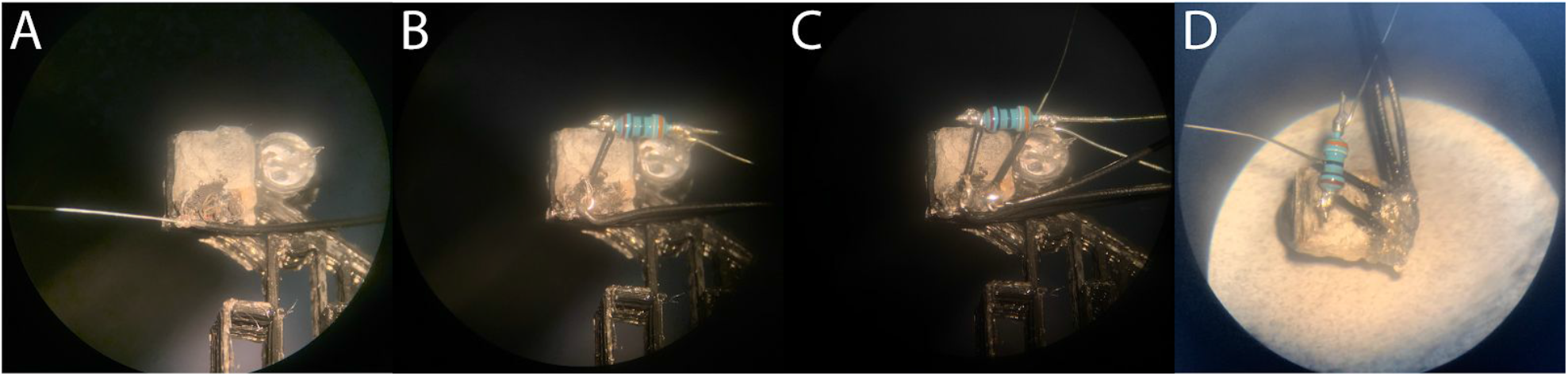

## Appendix 6: Shielding, bending, weaving, and stripping the wires

Close proximity of the wires and cameras to the headstage can introduce noise into the electrode channels. For this reason, we used aluminum tape to shield the electrical components of the headset. Cut a section of aluminum tape (Part #43) with a 5 mm wide strip running about 25 mm in length off of one side (A). Peel the backing from the strip, make two small cuts, and reinforce the joint with a small piece of tape or superglue to prevent tearing (A). Place the camera module at this joint, with the wires running on top of the strip (B). Wrap the strip around the three wires (C). Use a multimeter at this point to confirm that the aluminum is not electrically connected to any of the wires. Place the camera module in its proper position on a carrier and bend the wrapped wires at the proper locations so they feed towards the weaving lattice at the center of the carrier (D). Mark the location on the aluminum where the wires meet the first lattice crossbar on the carrier (shown by the red arrow in D). Remove the camera from the carrier, and peel back the aluminum strip to your marked location. Place the camera back on the carrier, and route the ends of the power and ground wires down through separate lattice windows of your choosing, and mark the location where they emerge from the bottom. It’s recommended to route the power wires in separate windows from the ground wires so they are well-separated when they are eventually soldered to the tethers as described in Appendix 9. Remove the camera from the carrier, and strip the ends of the power and ground wires to where you made your mark. Repeat this for each camera while it is in its proper position.

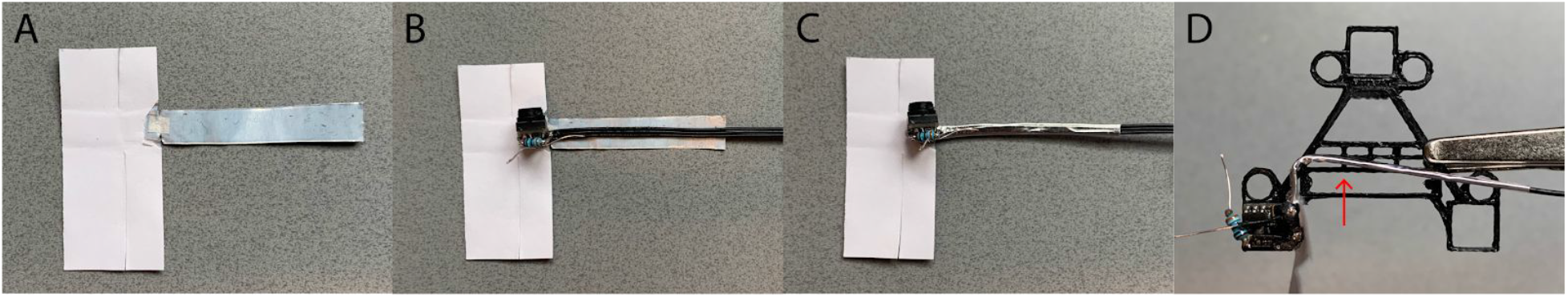

## Appendix 7: Cementing the cameras onto the carrier

Place the camera on the carrier with the power and ground wires in their proper windows, and weave the camera’s signal tether down through a lattice window, and back up, so it emerges upward from the center of the carrier (A). Slightly remove the camera from its holder, apply a small amount of superglue gel around the sides, and place it back in its holder, ensuring that the surface of the camera is perfectly flush with the bottom surface of the camera carrier (B). Once the glue has completely cured and the camera is solidified into place, peel off the paper from the unapplied aluminum tape, and wrap the camera module so it is as completely encased as possible, adding silicone glue or PTFE tape (Part #44) between the aluminum and camera components for insulation as needed, to ensure they don’t become electrically connected. Confirm this using a multimeter. Repeat these steps for the remaining cameras.

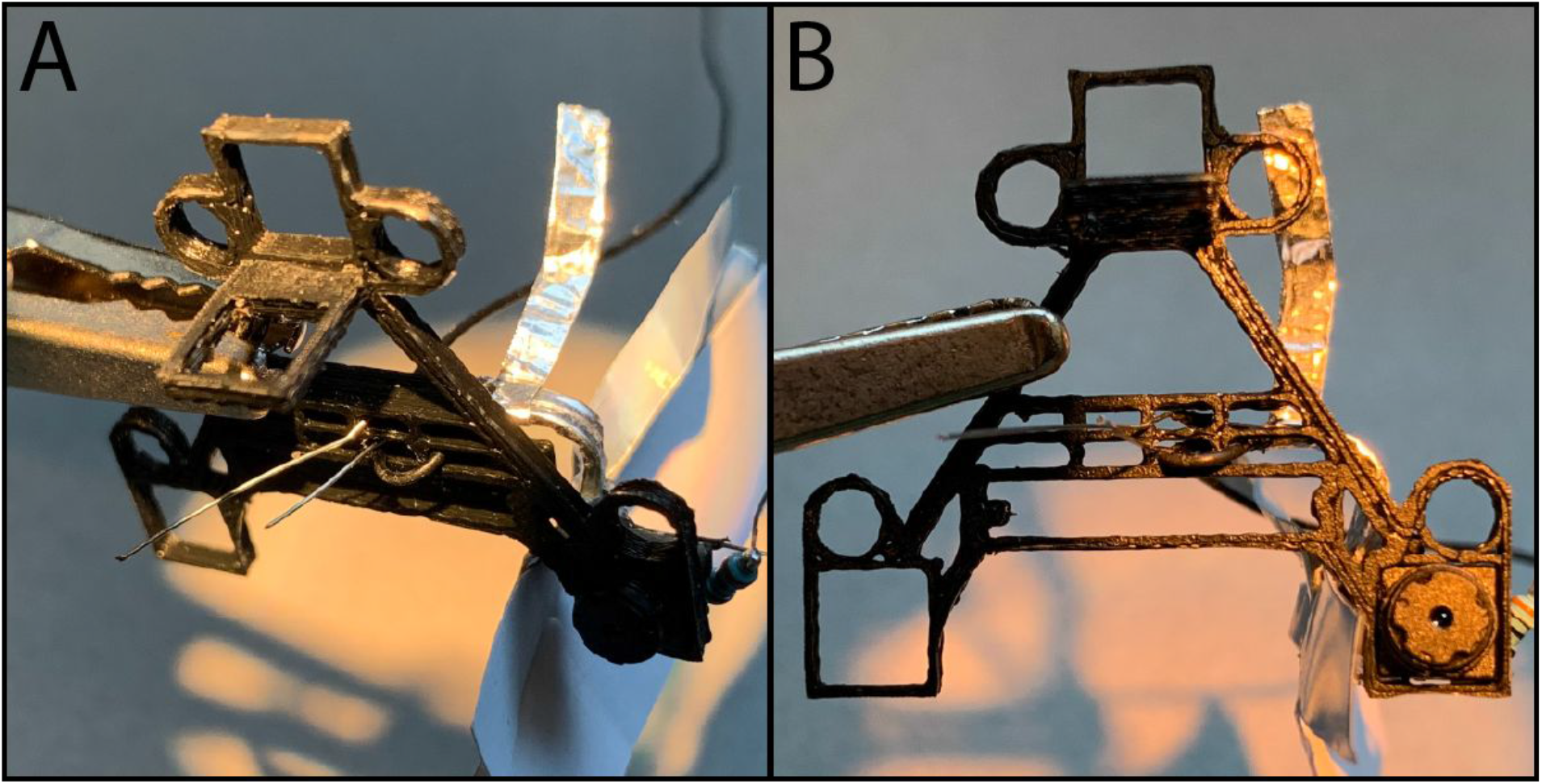

## Appendix 8: Adding the LEDs

Trim the power and ground terminals of an IR LED, and insert it into an LED receiver on the carrier (shown by the black arrow in A). Wrap the local power and ground wires from the camera module to the leads of the LED (shown by the red and black arrows in B). Use a multimeter to ensure that no undesired connections to the shielding exist, and then solder them in place for a stable and permanent connection. Repeat these steps for the remaining LEDs.

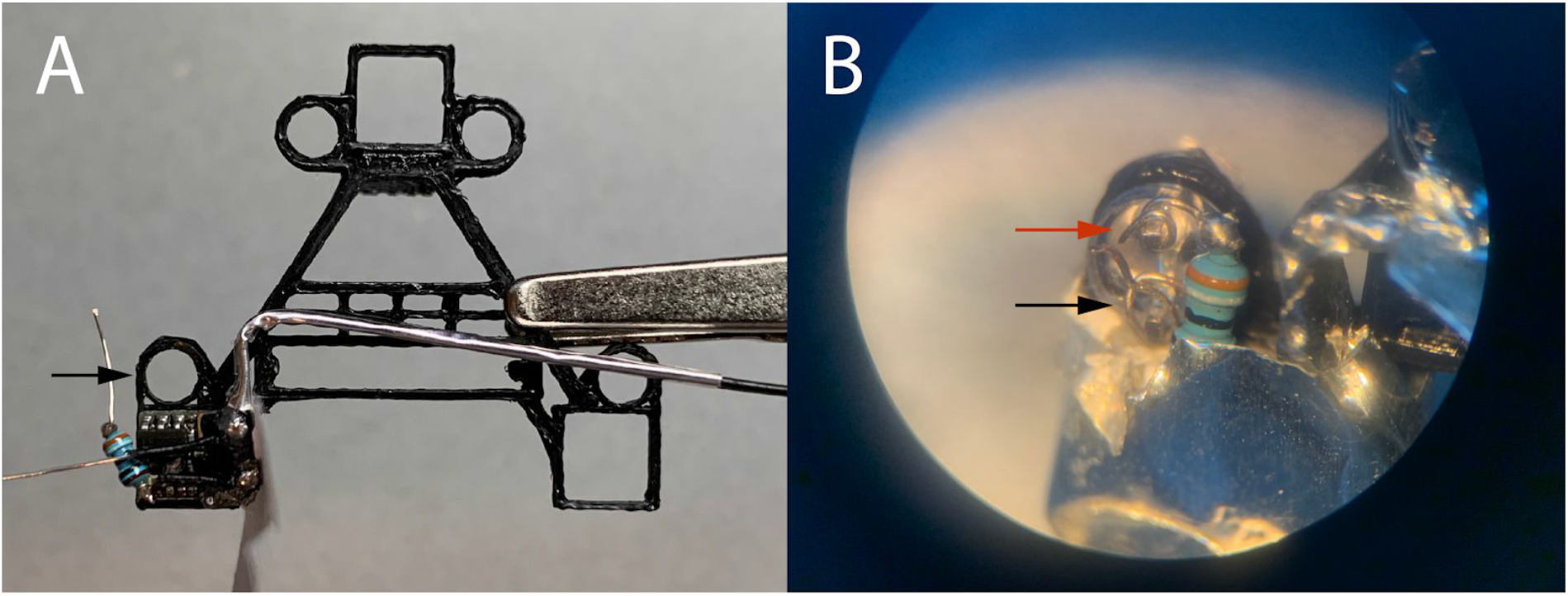

## Appendix 9: Soldering the power and ground tethers

With all the cameras cemented in place and their power and ground wires extending from the bottom of the carrier (A), prepare two wires cut to your desired tether length to serve as the common power and ground tethers. Strip the ends of the wire, and feed it down through the appropriate window for the power or ground. Twist the 4 power (or ground) wires around the respective tether wire (B). Ensure that there is no connection between the power and ground wires using a multimeter. At this point, you should power the cameras to test that they all produce a proper video signal before moving forward. Solder the wrapped wires for a permanent connection, and completely cover this bare wire and solder with insulation so they can be wrapped with aluminum shielding in the next step.

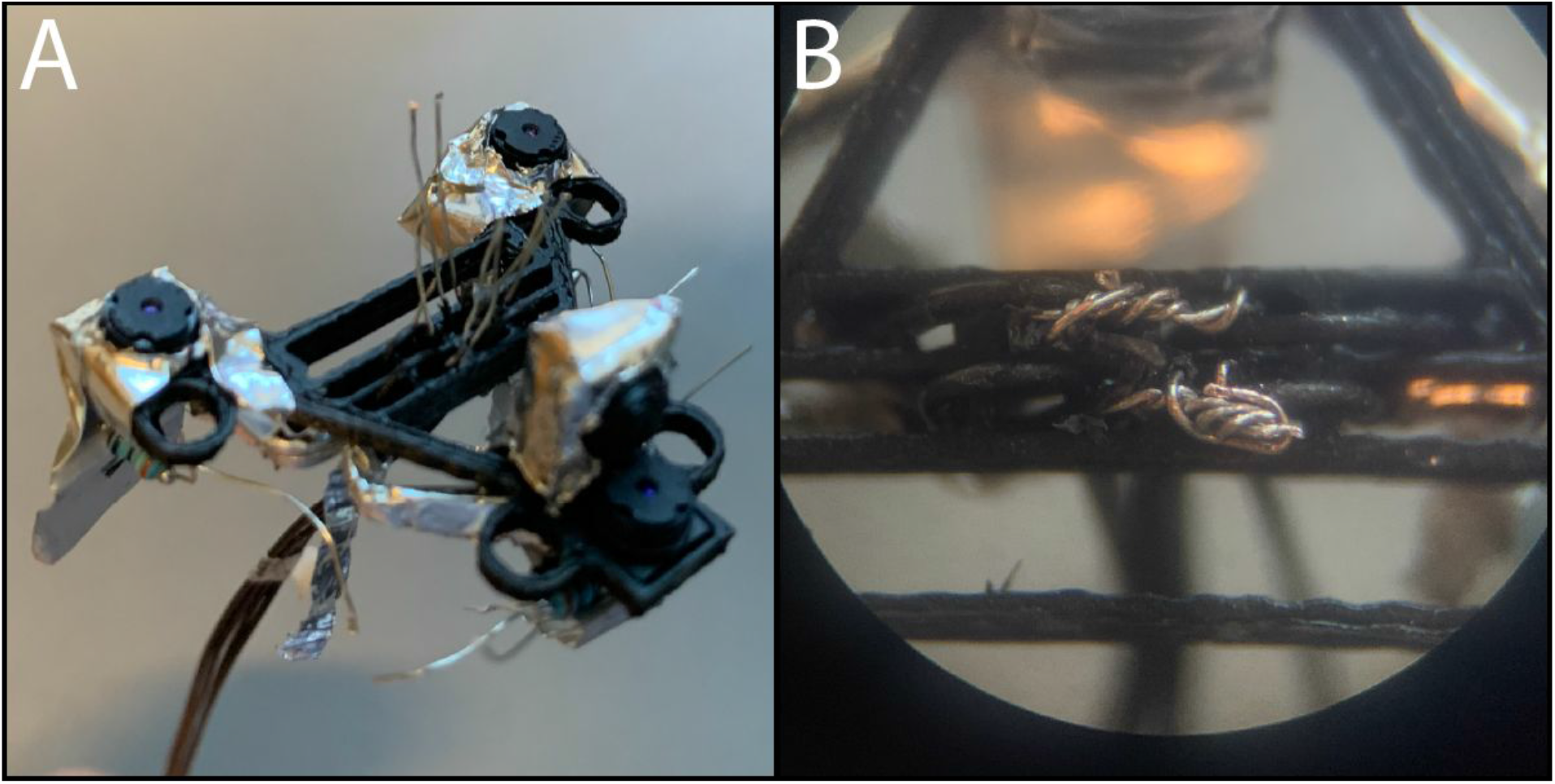

## Appendix 10: Shielding the weaving lattice and initial segment of the tethers

Cut a 75×15 mm piece of aluminum tape, with 2×1 mm notches on each side about 11 mm from the top (A). Peel and remove the backing from the top to the point of the notches (B). Replace the backing onto the aluminum, and fold the top portion as shown (B). For clarity, the next steps are shown with an empty carrier. Insert the top portion through the front opening, from the underside of the carrier (C). Remove the protective paper (D). Insert the remaining end through the front portion of the headstage window (E). Pull both sides upwards, so they surround the weaving lattice as shown (F). Press them together, so they encase the woven wires (G) and tethers (not shown). Trim and fold the remaining tape so the first 4cm of the tethers are completely encased, shown here on a prepared carrier (H). All shielding combined, as shown in (H), weighed 84 mg.

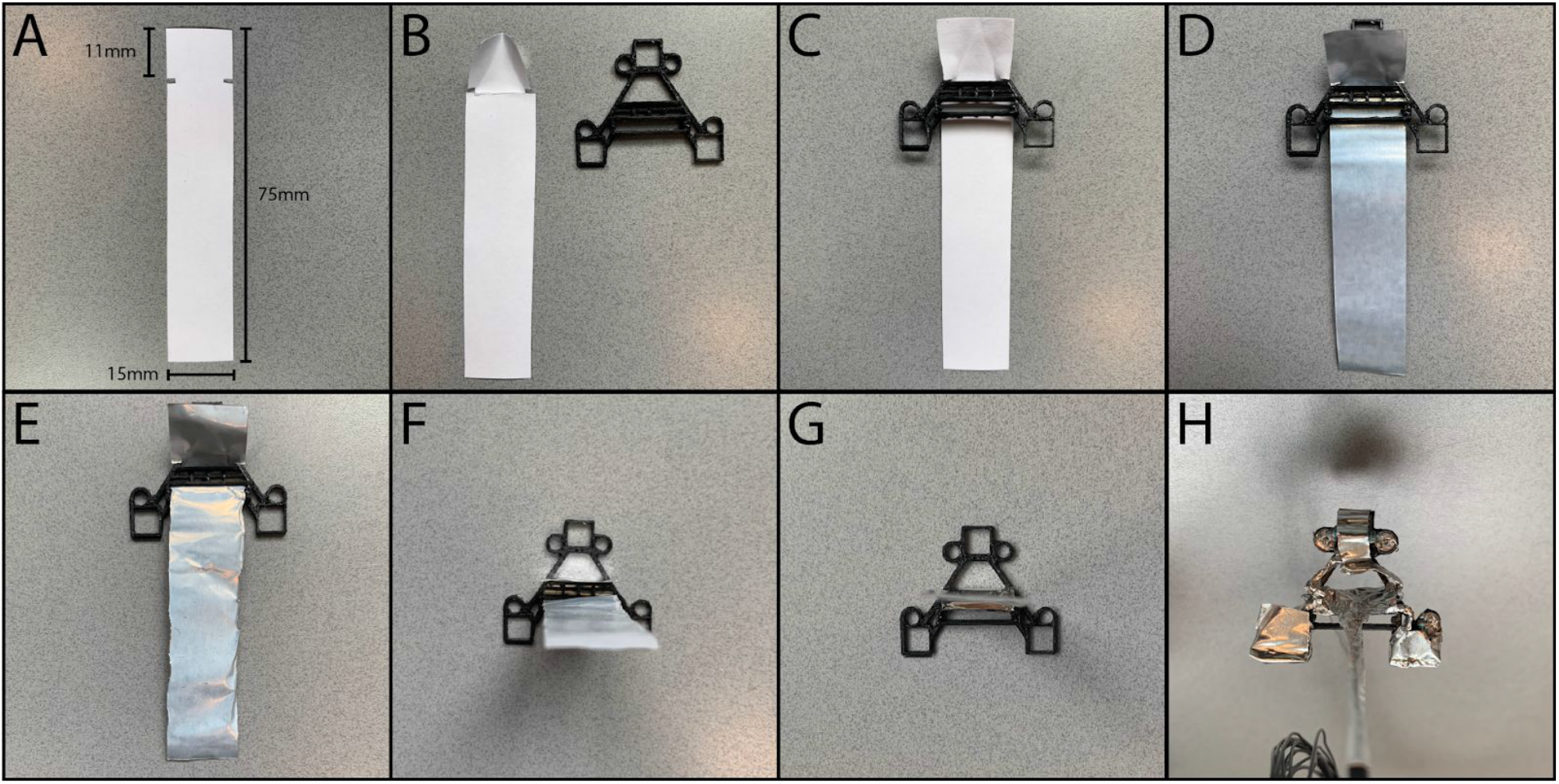

## Appendix 11: Attaching the headset to a headstage

Unplug a headstage, and slide it up through the back of the headstage window on the headset with the front of the headstage facing forward, as seen with an empty carrier as an example in A. The fit is very tight, so we recommend practicing this on an empty carrier first to get an idea of how much force you’ll need to use. Lower the headset until it reaches the bottom of the headstage as shown in B. Once placed on a headstage, the headset can be removed and reattached if necessary, but may cause wear and tear to the headset, and should be avoided.

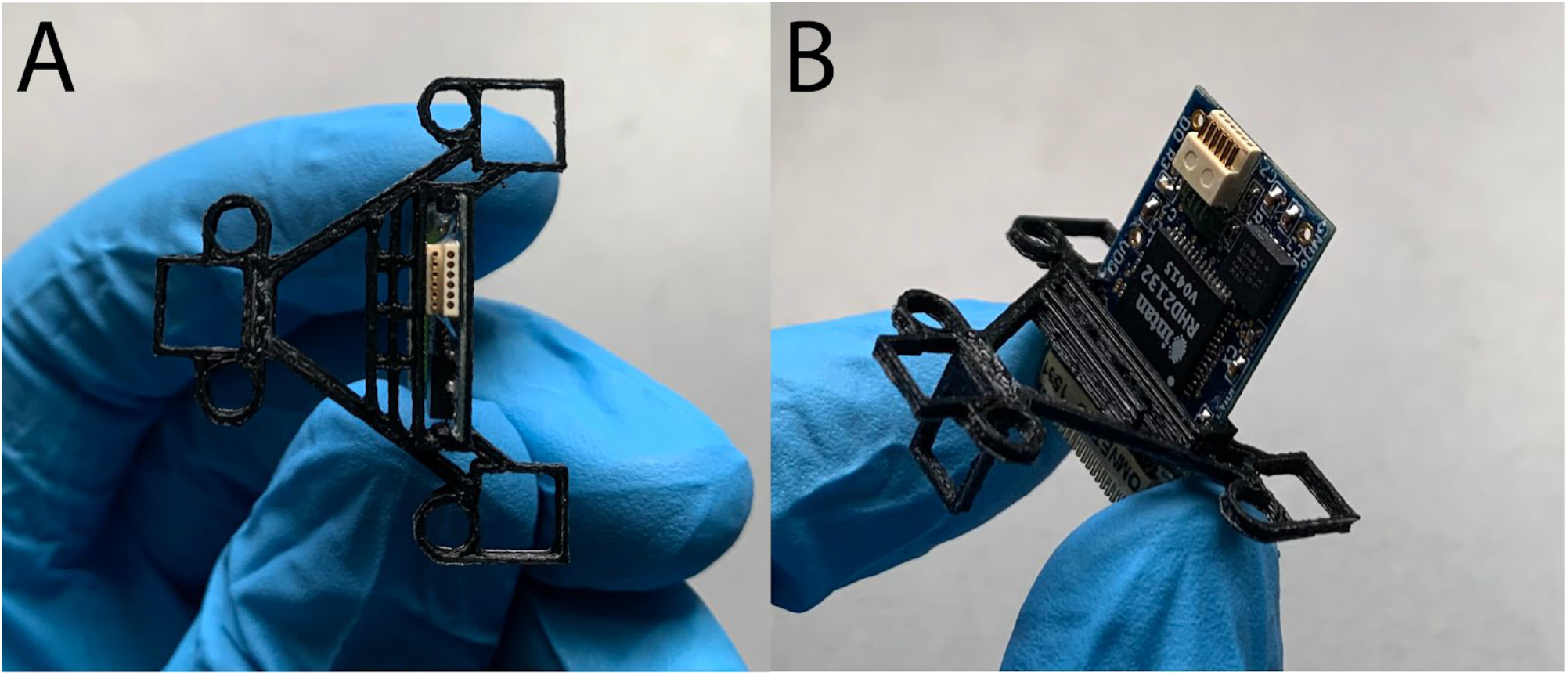

## Appendix 12: Grounding the shields

Prepare stripped wire to serve as the grounding connections for the shields. Create a small puncture in each of the aluminum shields of the headset, and weave the wire through them to create an electrical connection. Use a multimeter to confirm that all pieces of shielding have a low resistance connection to the remaining free end of the shielding wire. Use the multimeter once again to confirm that none of the shields have an electrical connection to any camera components at this point as well. Use silver conductive paint (Part #45) to adhere the shielding wire to the shielding tape at each connection point. Pin the free end of the shielding wire to either the GND or REF of the headstage with a large gold pin (Part #46), and coat it in silicone glue for a stable, yet removable connection. If pinning to REF, you’ll need to shorten the length of the pin, and pin from the front of the headstage so it can make a solid connection without hitting the back of the carrier.

## Notes

The authors state that they have no conflicts of interest.

### Competing Interest Statement

The authors have declared no competing interest.

